# Characterizing the Spatial Distribution of Dendritic RNA at Single Molecule Resolution

**DOI:** 10.1101/2024.07.18.602927

**Authors:** Jihoon Kim, Jean G. Rosario, Eric Mendoza, Da Kuang, Junhyong Kim

## Abstract

Neurons possess highly polarized morphology that require intricate molecular organization, partly facilitated by RNA localization. By localizing specific mRNA, neurons can modulate synaptic features through local translation and subsequent modification of protein concentrations in response to stimuli. The resulting activity-dependent modifications are essential for synaptic plasticity, and consequently, fundamental for learning and memory. Consequently, high-resolution characterization of the spatial distribution of dendritic transcripts and the spatial relationship across transcripts is critical for understanding the pathways and mechanisms underlying synaptic plasticity. In this study, we characterize the spatial distribution of six previously uncharacterized genes (*Adap2*, *Colec12*, *Dtx3L*, *Kif5c*, *Nsmf*, *Pde2a*) within the dendrites at a sub-micrometer scale, using single-molecule fluorescence in situ hybridization (smFISH). We found that spatial distributions of dendritically localized mRNA depended on both dendrite morphology and gene identity that cannot be recreated by diffusion alone, suggesting involvement of active mechanisms. Furthermore, our analysis reveals that dendritically localized mRNAs are likely co-transported and organized into clusters at larger spatial scales, indicating a more complex organization of mRNA within dendrites.

## INTRODUCTION

The neuron is a morphologically polarized cell with highly specialized compartments: the central soma, a singular axon, and several dendrites that extend from the soma. These specialized compartments are morphologically and functionally distinct from the soma and each other, with their molecular composition partly being established by the localization of RNA (Eberwine et al. 2002, Martin and Zukin 2006, Holt and Schuman 2013, Hutten et. al 2014, Buxbaum et al. 2015, Van Driesche and Martin 2018, Biever et al. 2019, Holt et al. 2019). mRNA localization is of particular interest in the dendrites where it allows for local and spatially restricted translation of protein at a rapid time scale (Martin and Zukin 2006, Hanus and Schuman 2013), in each separate dendrites and even subdomains within each dendrite (Barbour 2001, Sun et al. 2021). Additionally, RNA localization allows for molecular composition of dendritic subdomains to be modulated dynamically in response to neurotrophic factors to maintain or alter the local proteome (Kim TK et al. 2013, Zappulo et al. 2017, Fonkeu et al. 2019). This modulation is critical to key neuronal functions including synaptic plasticity and retrograde signaling (Sutton and Schuman 2006, Buxbaum et al. 2015, Donlin-Asp et al. 2017, Holt et al. 2019). Furthermore, perturbations of RNA distributions within the dendrites can significantly alter local protein distributions and synaptic function, which have been linked to neurodevelopmental disorders (van Spronsen and Hoogenraad 2010, Penzes et al. 2011, Swanger and Bassell 2013, Donlin-Asp et al. 2017).

While previous works using sequencing or microarray have cataloged the dendritically present and enriched genes (Zhong et al. 2006, Poon et al. 2006, Ainsley et al. 2014, Taliaferro et al. 2016, Middleton et al. 2019), only a few high-resolution single-molecule RNA imaging studies have examined the spatial distribution at a fine scale (Cajigas et al. 2012, Buxbaum et al. 2014, Chen et al. 2020). The identification and characterization of common patterns of mRNA distributions on a sub-dendrite level, and their relationships to the dendritic geometries remains elusive and is still lacking for most RNAs. Moreover, the spatial relationship between different RNA species remains largely unknown. As the currently characterized mechanisms of RNA localization are specific to sets of genes (Mayford et al. 1996, Eom et al. 2003, An et al. 2008, Lee et al. 2016, Taliaferro et al. 2016, Tushev et al. 2018, Arora et al. 2022), hinting at diverse regulation of transport across dendritic mRNA, identifying patterns of colocalization between groups of genes could characterize new mechanisms. Moreover, identifying patterns of colocalization could reveal new relationships between genes involved in various downstream neuronal processes.

In this study, we characterized the spatial distribution of RNA molecules within dendrites of neurons using single molecule fluorescence *in situ* hybridization (smFISH). We found distinct spatial distributions of dendritically localized mRNA, dependent on both dendrite morphology and gene identity, which cannot be recreated by diffusion alone, pointing to more active mechanisms involved in their distribution. Additionally, our analysis demonstrates that dendritically localized mRNAs are not only co-transported but are also organized into clusters at larger spatial scales, indicating a more complex organization of mRNA.

## RESULTS

### single molecule (sm) RNA FISH and Comparison to Transcriptomics

We used multiplexed single molecule fluorescence *in situ* hybridization (smFISH), which allows for robust quantification and high spatial resolution of individual mRNAs. Broadly, we targeted six dendritically localized genes across three functional classes for our study: *Nsmf* and *Pde2a* are annotated to have dendrite and neurite related function (neuronal), *Dtx3l* and *Colec12* share immune response function (immune response), and finally, *Kif5c* and *Adap2* are associated with broader cellular function (miscellaneous). Briefly, the function of these localized genes are: *Adap2* (ArfGAP with dual PH domains2; metal binding, GTPase activity), *Colec12*, (collectin sub-family member 12; binding/phagocytosis of bacteria), *Dtx3L* (deltex 3-like, E3 ubiquitin ligase; cell damage repair, interferon-mediated antiviral responses), *Kif5c* (kinesin family member 5c; involved in synaptic transmission, mediates dendritic trafficking of mRNAs), *Nsmf* (NMDA receptor synaptonuclear signaling and neuronal migration factor; triggers changes in the cytoarchitecture of dendrites and spine synapse processes), and *Pde2a* (photodiesterase 2A, cGMP-stimulated; modulates presynaptic short-term plasticity). These genes were reported to be not only localized to, but also differentially enriched and consistently expressed in the dendrites as noted in several previous studies (Zhong et al. 2006, Cajigas et al. 2012, Ainsley et al. 2014, Tushev et al. 2018, Middleton et al. 2019). This allows us to assess the relationship between a gene’s functional class and its spatial distribution in neuronal dendrites.

Spectral constraints of the fluorescent reporters limited us to five channels, two of which are reserved for the dendrites and nuclei. We identified dendrites via MAP2 staining that should exclude axons; however, it is feasible that some of the captured neurites were axons, but they represent only one neurite out of many in any individual neuron. Briefly, for each available color channel, two gene probes were assigned (Cy3: *Colec12*, *Pde2a*; mCherry: *Dtx3L*, *Adap2*; Cy5: *Kif5c*, *Nsmf*). We performed multiplexed smFISH, three genes at a time, for eight (2^3^) combinations of gene probes across the available three-color channels. Each stained neuron was imaged as a Z-Stack at 100X (**Figure 1**). Next, binary masks separating the neuronal compartments, the soma and dendrites, were generated per sample using SynD (Schmitz et al. 2011). Finally, the binary masks and the ZStacks were analyzed with Aro (Wu and Rifkin 2015) to identify the pixel coordinates of individual mRNA molecules (**Figure 1**; see **Methods** for details). We analyzed 410 total neurons across the 8 gene combinations (**Supplementary Table 1**). Before comparing spatial distributions, we removed neurons that had FISH spot counts below the 2.5^th^ percentile, indicative of poor staining, or greater than the 97.5^th^ percentile, indicative of glial contamination or nonspecific binding, for any single gene or summed across all three genes in either the soma or the dendrites. We removed 90 neurons across the eight probe combinations and used a total of 320 neurons for downstream analysis (**Supplementary Table 1**).

**Figure 1:**
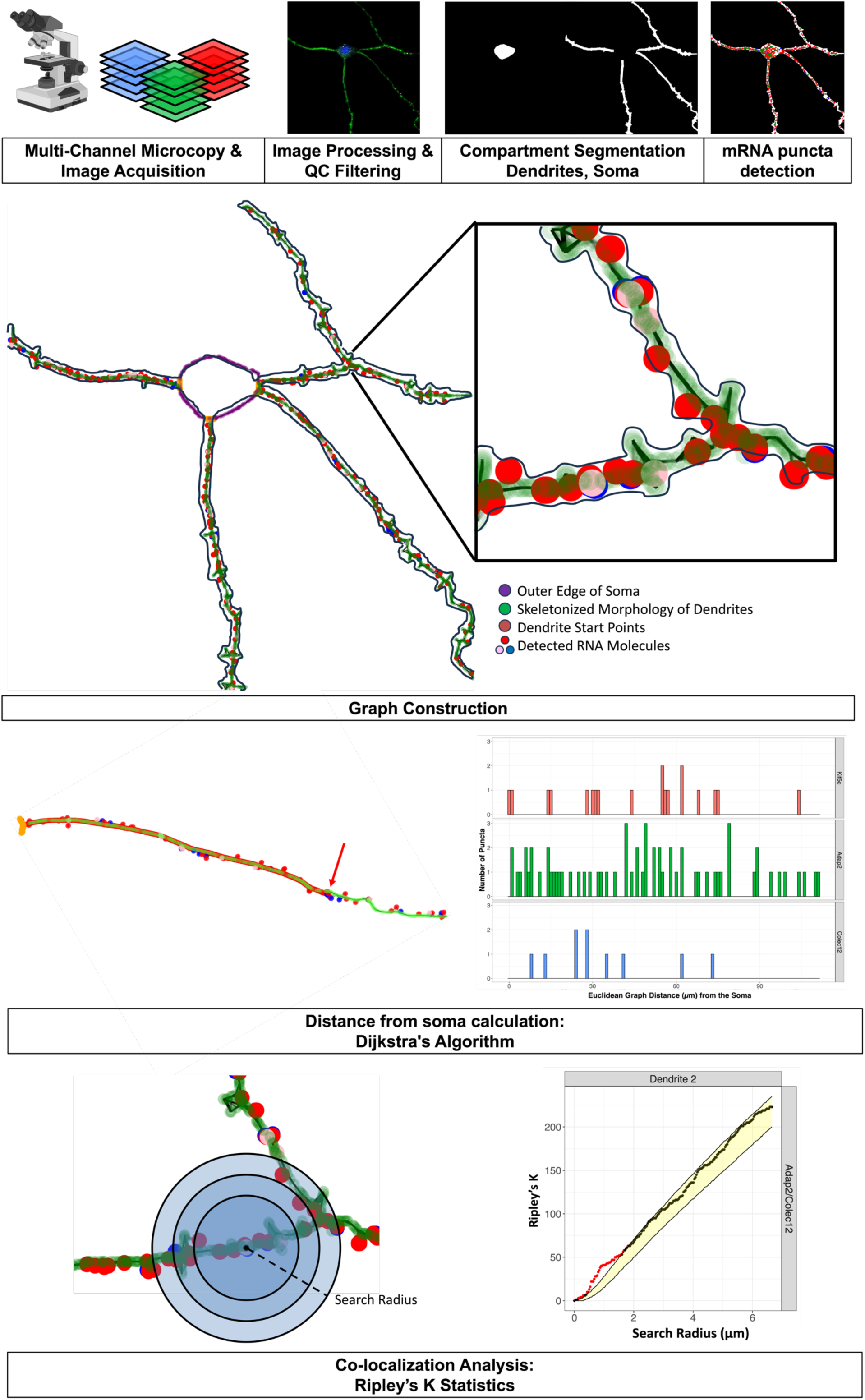
Outline of the characterization of mRNA spatial distribution within single neurons.

Comparing counts of each mRNA molecule per gene, we find that the distributions of the number of mRNAs detected per neuronal compartment (soma, dendrites in our study) in every sample across the triplet probe combinations were comparable. The respective mean counts per probe combination in each neuronal compartment were generally within one standard deviation from one another, though *Colec12-Adap2-Kif5c* probe combination, had lower counts for *Adap2*, and *Colec12-Dtx3L-Kif5c* had higher counts for all gene probed, *Colec12-Dtx3L-Nsmf* probe combination had higher counts for *Nsmf*, and *Pde2a-Adap2-Kif5c* had higher counts for *Pde2a*. Mean counts for *Adap2* in dendrites and *Pde2a* in soma were within one standard deviation across all probe combinations (**Supplementary Figure 1A**).

We found that the differences in the observed mRNA counts across probe combinations were partly explained by differences in the size distributions of the neurons stained. When we compared the neuronal compartment area, represented by the pixel counts of their respective binary masks, the distributions of areas were similarly comparable across probe combinations; the mean areas in pixels were within one standard deviation. There were two exceptions, *Colec12-Dtx3l-Kif5c* and *Colec12-Dtx3l-Nsmf* which had higher mean areas in both compartments (**Supplementary Figure 1B**). Generally, there was a positive correlation between the compartment area and detected number of mRNAs (**Supplementary Figure 1C**). Therefore, the larger compartmental areas for *Colec12-Dtx3l-Kif5c* and *Colec12-Dtx3l-Nsmf* probe combinations may explain the larger number of mRNAs observed for those probe combinations. However, for the case of *Colec12-Dtx3l-Kif5c*, which had a similar distribution of compartment areas to the other probe combinations, the low *Adap2* counts may be better explained by low quality staining or non-optimal imaging conditions. This decrease seems to be isolated to the probe targeting *Adap2*, as the remaining mRNAs (*Colec12*, *Kif5c*) were detected in similar quantities to the other probe combinations. Furthermore, there was no significant interaction effect, such as overcrowding or compensation, across the different combinations of gene probes or color channels used (data not shown). These similarities in count distributions indicate comparable expression levels across probe combinations.

### Comparison of mRNA Counts across Compartments

We found that the absolute number of mRNAs detected in the dendrites were higher than that in the soma, except for *Pde2a*, which was expressed in comparable quantities (**Supplementary Figure 1A**). When we compare the area of each compartment to the number of mRNAs detected within, we found that the number of mRNAs increased linearly with area, so the differences in the observed counts across compartments were due to differences in their size, specifically the area of the soma being smaller than that of the total area of dendrites (**Supplementary Figure 1B, C**). Therefore, to compare expression level of the genes more accurately across compartments, we compared their mRNA densities, the number of absolute counts divided by the pixel area of the compartment binary mask.

Generally, mRNA density in the soma is higher than that in the dendrites for all genes studied, though this disparity may be somewhat exaggerated by biases in estimating the compartment areas of the dendrites, due to their complex geometries (**Figure 2A**). Comparing mRNA densities per gene, we find that the distributions of the number of mRNAs detected per neuronal compartment in every sample across the triplet probe combinations were comparable; their respective means demonstrated similar trends to those observed across mean puncta counts discussed above (**Figure 2A, Supplementary Figure 1A**).As the differences observed across probe combinations likely represent small experimental variabilities and natural biological variation in neuronal culture, samples were combined across probe combinations for subsequent analysis. Regardless, we kept track of the role of these probe combinations as possible confounders in the downstream analysis, but did not find a specific effect, as discussed below.

**Figure 2:**
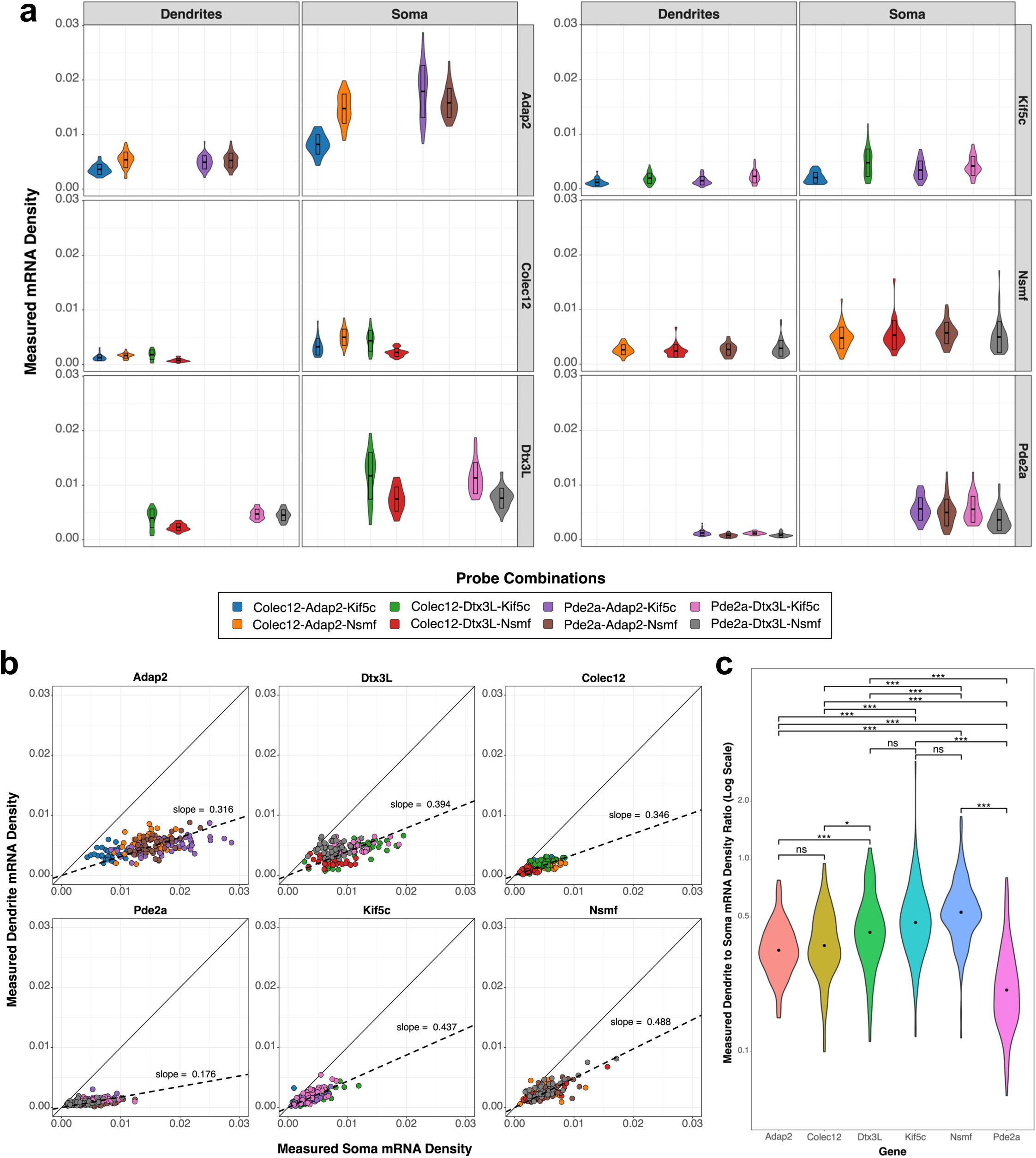
A comparison of gene-specific mRNA densities across probe combinations and neuronal compartments. **A)** A violin plot of mRNA densities (number of detected mRNAs per pixel area of neuronal compartment) across the triplet probe combinations. Probe combinations are shown in different colors. **B)** A scatter plot of dendrite mRNA density vs. soma mRNA density. The dashed line represents the linear regression line of best fit. **C)** A violin plot of compartmental density ratio across genes. Pairwise Kolmogorov–Smirnov test was performed. Each asterisk represents a distinct level of statistical significance: one asterisk indicates adjusted *p*-value < 0.05, two asterisks denote adjusted *p*-value < 0.01, and three asterisks represent adjusted *p*-value < 0.001.

When comparing mRNA densities across neuronal compartments, generally, an increase in mRNA density in one compartment was observed alongside an increase in mRNA density in the other compartment for all genes, except *Pde2a*. For *Pde2a*, while mRNA density in the dendrites was relatively consistent across the cells, the density in the soma were more variable across the cells (**Figure 2A**, **Supplementary Table 2, 3**). Additionally, the ratio of compartmental density (mRNA density in dendrite divided by mRNA density in soma) for each gene was consistent across neurons regardless of probe combinations (**Figure 2B**). We found significant differences in the dendrite to soma mRNA density ratios of the genes by pairwise Kolmogorov– Smirnov test (**Figure 2C**). Briefly, *Adap2* and *Colec12* share similar distribution of ratios; *Dtx3L*, *Kif5c* and *Nsmf* also share similar distribution of ratios, with *Pde2a* having a unique distribution of ratios. This gene group specificity is also supported by a pairwise Mann-Whitney U Test, where the mean dendrite to soma mRNA density ratios were statistically different across genes, except for *Colec12* and *Adap2*. Taken together, different genes seem to have different density relationships between the soma and dendrites, which points to a level of regulation of localization and not simply passive diffusion that is driving the spatial distribution of specific mRNA molecules in neuronal dendrites. The ratio of mRNAs’ density in the soma and dendrites likely reflects biophysical parameters determined by characteristics of the mRNA and the transport process. The differences in the ratio for some of the genes potentially suggest differences in transport regulation.

### Spatial Distribution of mRNAs along the Dendrite

To characterize the spatial distribution of the mRNAs within dendrites, we calculated each mRNA’s distance from the edge of the soma (**Figure 1**). The complex morphology of dendrites poses a significant challenge for characterizing accurate position of individual mRNA molecules. Given the non-linear and complex branching patterns of the dendrites and the irregular shape of the soma, conventional Euclidean distances do not accurately represent a mRNA molecule’s distance from the soma. To accurately compute distances from the soma, we constructed undirected graphs to approximate the geometry of the cell, then used Dijkstra’s Algorithm to calculate graph distances to approximate distances along dendrite geometry (**Methods**). We used these graph distances to characterize the density of mRNAs as a function of the distance from the soma (**Figure 1**).

The compiled distributions of computed mRNA distances appear similar across all genes probed. From the compiled distribution, it is evident that the number of detected mRNAs broadly decreases with distance from the soma for all genes probed in accordance with previous publications (Cajigas et al. 2012, Buxbaum et al. 2014, Chen et al. 2020) (**Figure 3A**). Additionally, a pairwise quantile-quantile plot of the compiled mRNA distance distributions per gene shows that distributions of mRNA densities are similar from the edge of the soma to about 100 - 150 μm away from the soma for all gene pairs; the primary differences between distribution lie towards the distal ends of the dendrites (**Supplemental Figure 2A**). Specifically, the genes tended to show density differences in the range of 160 - 190 μm, likely due to the variations in the size of the neurons stained. In fact, some genes (*Adap2*, *Kif5c*, and *Pde2a)* had greater numbers of dendrite masks with very large areas (suggesting longer dendrites) than others suggesting the raw distribution is compounded with the size of the imaged neurons (**Supplemental Figure 1C**). The mean mRNA distance from the soma, while generally statistically different (Mann-Whitney U Test), only differed from each other by 1 - 2 μm, except for *Pde2a*, whose mean mRNA distance was smaller by about 5 μm (**Supplementary Table 4**). This difference in mean mRNA distance could indicate that *Pde2a* may be actively localized closer to the soma. Collectively, the distribution of mRNA molecules along the dendrites by raw distance is similar across the genes we probed up to 150um, suggesting that the effect of gene identity may be limited in the spatial localization of these six genes in the dendrites for relatively proximal distributions.

**Figure 3:**
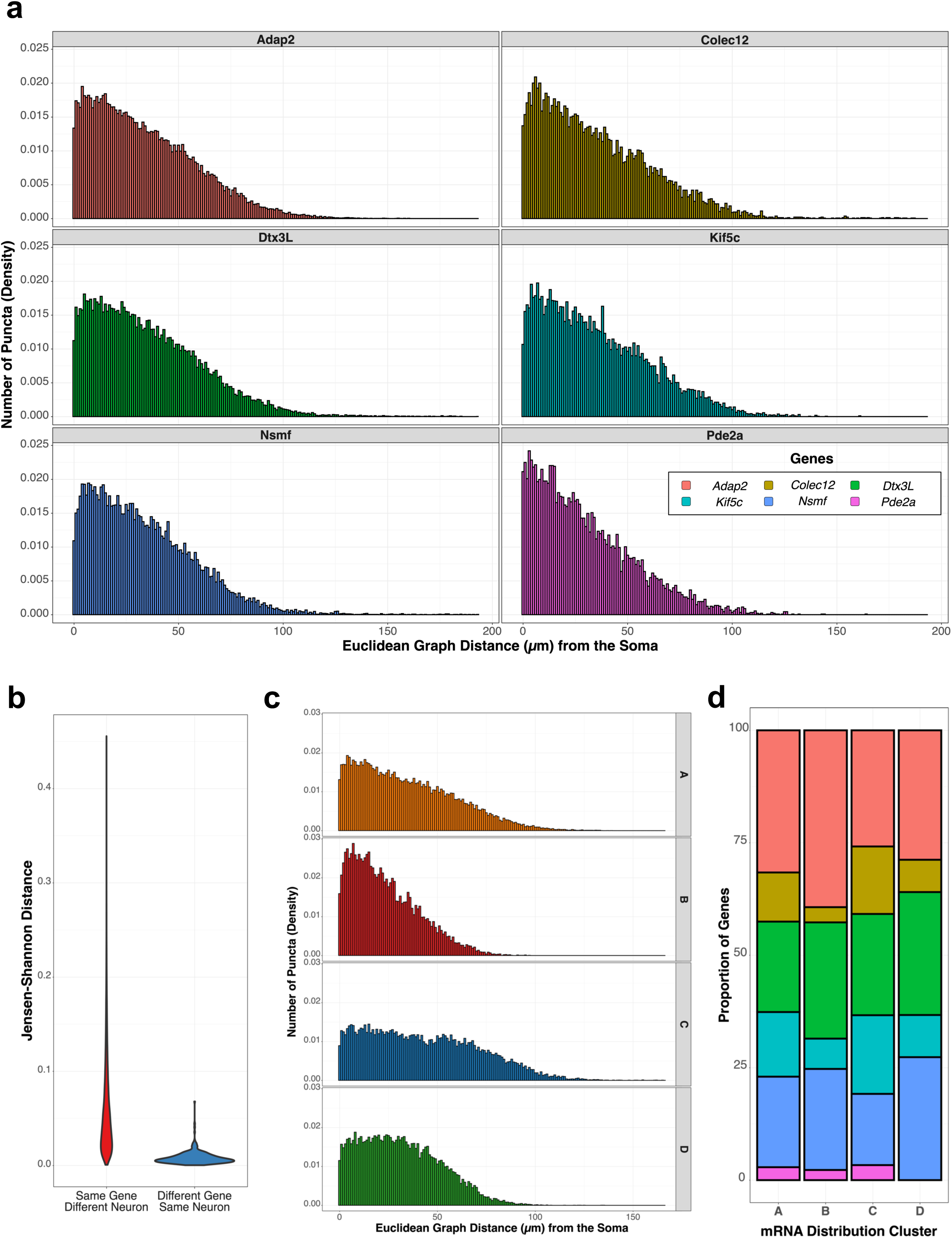
Characterizing the spatial distribution of dendritically localized mRNAs. **A)** Histograms of graph distances from the soma of dendritically localized mRNAs plotted per gene. Binwidth is 1 µm. **B)** Violin plot of Jensen-Shannon distances of mRNA distance distribution comparing two different neurons probed for the same gene (red) and mRNA distance distributions comparing different genes probed within the same neuron (blue). Distance values closer to 0 represent a higher degree of similarity between distributions. **C)** Histogram of mRNAs’ distances from the soma specific to each spatial patterning cluster. **D)** Bar graph representing the proportion of distance distributions from each gene that was assigned to each spatial pattern cluster. Genes are represented in separate colors.

While the general trend of decreasing density with distance is shared across the genes, at a more detailed scale, some features were different between the genes. Notably, the distributions of mRNA distances were not monotonically decreasing (**Figure 3A**); the density of mRNAs detected increases from the perimeter of the soma to about 5 μm away from the soma before decreasing, indicating a zone of mRNA depletion for certain genes. This pattern may imply a regulatory mechanism for mRNA localization, though it may also be an artifact from our image compartment segmentation. Since our morphological stain primarily targets dendrites (MAP2), we dilated the binary mask of the soma, which was not as brightly stained, in case the soma-dendrite boundary delineation caused the initial drop in densities. However, on average, only 3.5% of the mRNAs were affected in each neuron by this possible assignment discrepancies (data not shown). Recalculating the mRNAs’ distances from the soma with the un-dilated masks did not result in a monotonically decreasing distribution, when compiled (data not shown).

The initial increase in mRNA density around 4 - 5 μm contrasts with the previous model/observation reported (Fonkeu et al. 2019), where they had reported that spatial mRNA profile can be described by an exponential function whose exponent is a function of the half-life, diffusion, and transport velocity; the shapes of our compiled distributions of computed mRNA distances do not follow the shape noted in Figure 2 of Fonkeu et al. 2019. One explanation is that the previous exponential model does not account for dendrite length or morphology, which affect the spatial distribution of the mRNAs within dendrites, as we discuss below. Alternatively, the initial portion of the dendrite may act as a filter for the selection of dendrite-specific cargoes as have been reported in axons (Song et al. 2009, Watanabe et al. 2012, Petersen et al. 2014).

### Characterizing Patterns of mRNA Distribution

We identified common patterns of mRNAs’ distance distributions, representing similarities in their locations along the dendrites. Our dataset consists of 320 neurons, each multiplexed for three genes, resulting in 960 gene-neuron combinations. Ignoring the gene-neuron identities, we characterized the similarities between the distance distributions, or common spatial patterns across the neuron-gene pairs, using Jensen-Shannon distances (JSD). To accurately calculate the JSD between any two distributions, we first filtered for distributions with at least 100 spots detected, resulting in 537 mRNA distance distributions. We then calculated the pairwise JSD across the distributions and used the resulting distance matrix to perform hierarchical clustering. From our clustering analysis, we identified four separable clusters consisting of distributions from at least 10 neuron-gene pairs (**Supplementary Figure 2B**). The clusters represent unique patterns of mRNAs’ localization along the dendrites, regardless of any probe combination or gene **(Supplementary Figure 2D)**. There was no strong association between the proportions of staining combination and the four pattern classes demonstrating that our probe combination does not introduce a strong bias in spatial patterning. Genes were found in similar proportions across the identified clusters (**Figure 3D**), but, clusters and gene identities were not completely independent (Pearson’s Chi Square Test), supporting the previous report that gene identity influences mRNA’s spatial location (Cajigas et al. 2012). Nevertheless, mRNA distance distributions comparing different genes probed within the same neuron were more similar than mRNA distance distributions comparing the same genes probed between different neurons (**Figure 3B**). Our analysis suggests that dendritic morphology may be a stronger factor in determining the spatial distribution of localized genes.

We then aggregated the mRNA distance distributions per cluster to visualize the spatial patterning represented by the clusters (**Figure 3C**). Previously, Cajigas et al. (2012) found three large classes of spatial patterning of mRNAs along the proximal-distal dendritic axis that differ in the rate of decline in the number of particles along the dendrites. Our clusters match two of the clusters Cajigas et al. reported. Briefly, our clusters **A** and **B** represent the class where the mRNA density along the dendrite decreases generally monotonically (*generally monotonic-decay*), while clusters **C** and **D** represent the class where mRNA densities are maintained before decreasing (*plateau-decay*) as the distance from the soma increases (**Figure 3C**). The last class in Cajigas et al., representing an early and sharp decline of mRNA molecules, were not found in our study as the genes analyzed here were those with high dendritic expression (Cajigas et al. 2012). Our clusters **A** and **B** within the *generally monotonic-decay* class differ in the rate of decay, where cluster **A** decays more slowly than cluster **B**, resulting in a longer tail for cluster **A**. Our clusters **C** and **D** within the *plateau-decay* class differ in where decay begins after the plateau; cluster **C** has a longer plateau before decay and a corresponding longer tail (**Figure 3C**). We hypothesized that the differences in the maximum distance a mRNA molecule was found may be driven by the distributions in the physical size of dendrites/neurons. Specifically, as longer dendrites are rarer, the physical size of the dendrites would limit the number of mRNAs found at larger lengths away from the soma. This is supported by the distributions of dendrite area per cluster, a proxy for the dendrite length assuming consistent width across individual cells. The areas of the longer tailed distributions (cluster **A** and **C**) were larger than their shorter tail counterparts (clusters **B** and **D**) (**Supplementary Figure 2C**). Accordingly, the observed differences in the tails of the distributions within each class, suggests that the geometry of the dendrites, specifically their length, within which the mRNAs were detected, may be a determining factor in the observed spatial distribution of the mRNAs. It should also be noted that we cannot easily differentiated between true geometry of the cell and the geometry determined by the limited field of view of our imaging system.

The spatial patterns characterized do not follow the shapes of RNA distributions predicted by exponential function reported in Fonkeu et al. 2019, as discussed above. However, our spatial patterns seem to correspond to the three fundamental protein distributions described in Fonkeu et al. 2019: 1) monotonically decreasing density as a function of distance from the soma, corresponding to cluster **A**, 2) a plateau in density outside the soma followed by an exponential decrease, corresponding to cluster **C** and **D**, and 3) an increasing density at proximal distances from the soma followed by a peak and decrease, corresponding to cluster **B**. The similarities between the observed spatial patterns to the described protein distributions may be explained by local, spatially restricted translation (Martin and Zukin 2006, Hanus and Schuman 2013).

### Accounting for Length Dependence in Puncta Distributions

As we found that dendritic morphology, especially length, may affect the spatial localization of the mRNAs, we re-characterized the spatial distribution of the mRNA molecules within individual dendrites. We manually segmented well-separated, linear, single dendrite binary masks from our dendrites’ binary masks. We were careful to select single dendrites that had minimal bumps, no branching, and no overlap with other dendrites to minimize ambiguity **(<u>Methods</u>)**. From 320 neurons analyzed, 230 neurons had at least one clearly separable dendrite, from which we curated 411 single dendrite masks. For each single dendrite mask, we calculated the *raw graph distance* from the soma for each detected mRNA molecule, as before. Generally, the compiled distribution of mRNAs’ raw distances at the single dendrite level was similar to the compiled distribution raw distances at the neuron level. There was a broad decrease in mRNA density with distance from the soma and the density of mRNA detected increases from the perimeter of the soma to about 5 μm away from the soma before decreasing for all genes probed (**Supplementary Figure 3A**). As the probability of branching increases with length, our dataset of a single dendrites is a subsample from all single dendrites in the neurons, biased for shorter dendrites. Therefore, the compiled distribution at the single dendrite level is also a biased subsample of the compiled distribution at the neuron level, missing longer distances, as visualized in the shorter tail and the shift in the quantile-quantile plot **(Supplementary Figure 3A, Supplementary Figure 3B)**. To account for the differences in lengths across the single dendrites analyzed, we measured the length of each single dendrite mask, and computed the *normalized* distance from the soma by dividing the mRNAs’ *raw graph distances* by the dendrite length **(<u>Methods</u>)**. When the mRNAs’ *normalized* distances were compiled, we found that mRNA density decreased much more slowly, generally persistent at high levels throughout the entire length of the dendrite, for all genes probed (**Figure 4A**).

**Figure 4:**
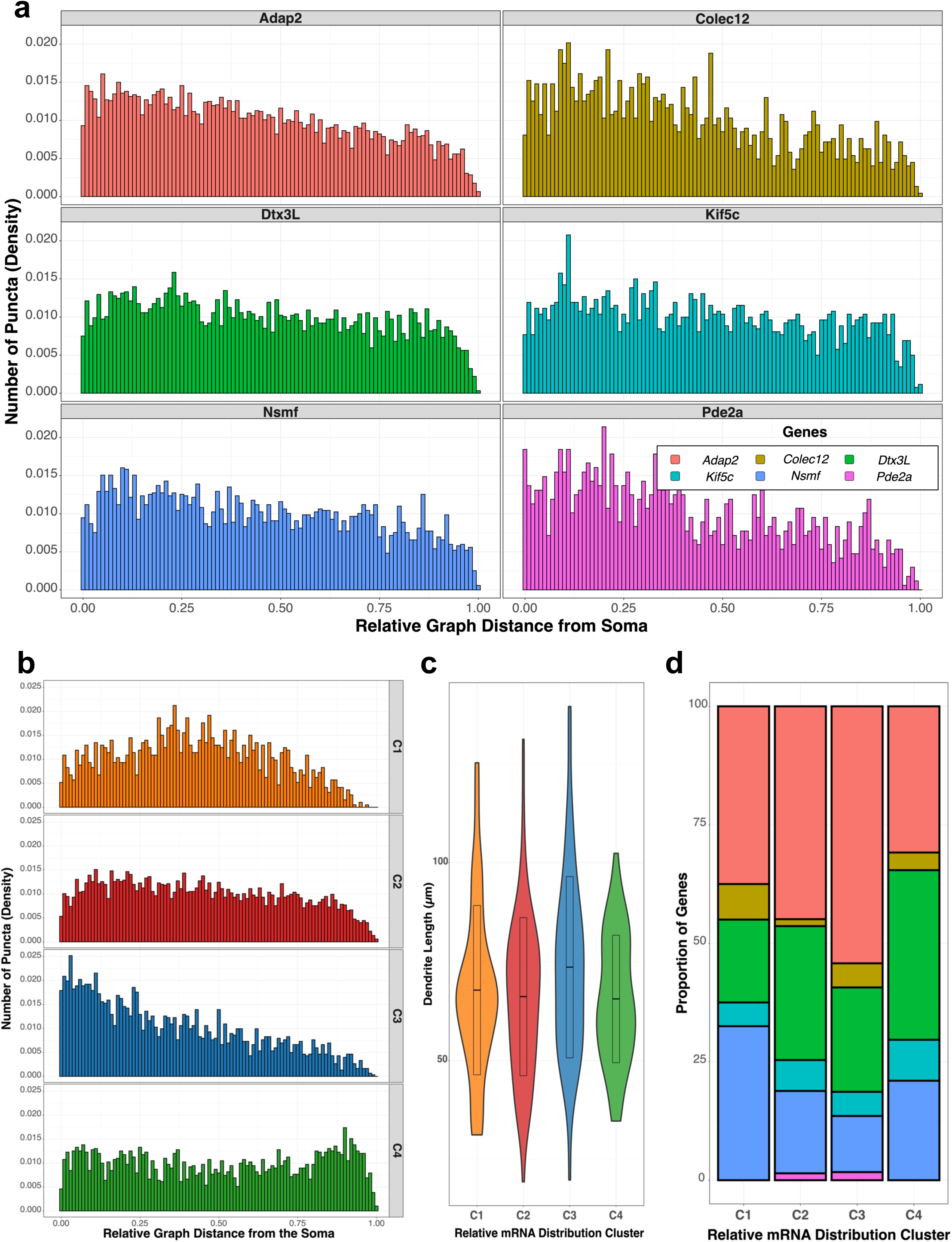
Characterizing the spatial distribution of mRNAs within single dendrites to account for dendrite length. **A)** Histograms of *normalized* graph distances from the soma of dendritically localized mRNAs plotted per gene. **B)** Histogram of mRNAs’ *normalized* distances from the soma specific to each spatial patterning cluster. **C)** Violin plot of dendrite lengths of dendrites assigned to each spatial patterning cluster identified. **D)** Bar graph representing the proportion of distance distributions from each gene that was assigned to each spatial pattern cluster. Genes are represented in separate colors.

Next, we identified common patterns of mRNAs’ *normalized* distance distributions from our single dendrite analysis. We compiled 411 (dendrites) X 3 (genes) = 1,233 gene-dendrite *normalized* distance distribution across 230 neurons; that is, each mRNA *normalized* distance distribution represents one of the 1,233 neuron-dendrite-gene combinations. We first filtered for distributions with at least 30 mRNAs detected, resulting in 320 mRNA distance distributions, from 163 neurons. We selected 30 mRNAs per dendrite as the minimum threshold to keep in line with our previous neuron level analysis, where we chose 100 mRNAs per neuron dendritic arbor, and we estimated that each neuron had at least three to four dendrites. Similar to our neuron level analysis, we calculated the pairwise JSD across the *normalized* distance distributions and used the resulting distance matrix to perform hierarchical clustering. From our clustering analysis, we identified four separable clusters consisting of distributions from at least 10 neuron-dendrite-gene combinations, which are distinct from the previous four clusters identified in the neuron-gene pair analysis (**Supplementary Figure 3C**). These clusters of *normalized* distance distributions represent unique patterns of mRNAs’ *relative positions* along the dendrites.

Briefly, we classify the clusters of *normalized* distance distributions as: (**C1)** a symmetric distribution of mRNAs with the highest mRNA density at the center of the proximal-distal axis of the dendrite; (**C2)** generally uniform distribution of mRNA density with a slight decrease towards the distal end of the dendrite; (**C3)** monotonically decreasing along the length of the dendrite; (**C4)** generally uniform distribution with the highest mRNA density at the distal end (**Figure 4B**). We observe that cluster-specific patterns are reflected on an individual neuron-dendrite-gene combination, despite a high degree of variability, indicating that the patterns are not a product of aggregating the distributions (**Supplementary Figure 3D**). Additionally, the distributions of dendrite lengths were comparable across clusters, confirming that the spatial patterns represented by these clusters were not dependent on the particular mixture of lengths of the dendrite (**Figure 4C**).

These patterns of mRNAs’ relative positions were not specific to any probe combinations (**Supplementary Figure 3E**) nor were dependent on the numbers of detected mRNA molecules within each dendrite (**Supplementary Figure 3F**). Similar to our previous neuron level clustering analysis, each of the newly identified clusters contained *normalized* distance distributions from every gene, except for *Pde2a*, which was only found within Clusters **C2** and **C3**, which is due to only three single dendrites containing more than 30 detected mRNA molecules of *Pde2a.* Furthermore, unlike the neuron-level analysis, there was no statistically significant relationship between cluster membership and gene identities (Pearson’s Chi Square Test). However, certain genes do appear enriched in specific clusters. Notably, *Adap2* was found in higher proportion in **C3**, *Dtx3L* was found in higher proportion in **C4**, and *Nsmf* was found in higher proportion in **C1** (**Figure 4D**). While we cannot discount the possibility that gene identity might still have a small effect, as observed on the neuron-level analysis, our analysis suggests that dendrite length has a stronger correlation to the spatial distribution of localized genes.

In cluster **C1** the maximum mRNA densities were at the relative center of the dendrite, while for C4 the maximum densities were found at the relative distal end of the dendrite, respectively. This is in stark contrast to the patterns identified in both our previous neuron level analysis, and Cajigas et al, where the maximum mRNA density was consistently found at the proximal end of the dendrites nearest to the perimeter of the soma. This difference highlights the sensitivity of the spatial distribution of dendritically localized mRNAs to the length of the dendrite. This may explain why previous studies, which did not normalize for the length of the dendrites analyzed, reported a sharp decline of mRNA densities for genes that may have functionality throughout the entire dendrites, such as β-actin (Buxbaum et al. 2014). More importantly, our analysis reveals four distinct classes of dendrites differentiated by the locations of peak mRNA density; these locations may represent functionally important regions within the dendrite, such as regions of high spine or synaptic density. Previous studies, which did not normalize for morphological features of the dendrites analyzed, only reported a single class, which showed an initial high density of mRNA molecules at the base of dendrites that decreased over the length of the dendrite (Cajigas et al. 2012, Buxbaum et al. 2014, Fonkeu et al. 2019, Chen et al. 2020).

### Identifying Morphological Features of Dendrites that affect mRNA Distributions

As both dendrite morphology and gene identity were found to affect the localized mRNAs’ spatial distributions, we sought to build a predictive model of localization distribution classes (i.e., the four clusters) using features that might influence spatial distributions. To this end, we compiled the binary masks, original Z stacks, as well as the maximum projections of the single dendrites used for the clustering analysis and prepared one-hot vectors representing the gene identity for each data instance.

Initially, we built a model to predict cluster labels from gene identity features using a random forest algorithm. Given the limited and unbalanced nature of the dataset with 40 instances for cluster C1, 138 for cluster C2, 59 for label C3, and 81 for cluster C4, we implemented 5-fold cross-validation. In each fold, clusters 1 and 3 were oversampled to 50 for training, and the class weight in the random forest was inversely adjusted to class frequencies. We ended with a weak classifier with a mean accuracy of 0.29 ± 0.03 on training and 0.26 ± 0.02 on testing, with F1-scores of 0.22 ± 0.02 and 0.19 ± 0.02, respectively.

We next trained a convolutional neural network using three-channel images (binary-mask, max-project, z-stack) combined with the one-hot vector of gene identity, aiming to predict cluster distribution labels (**Figure 5A**). With data augmentations, balanced sampling, and regularization (weight decay = 10^-5^, dropout = 0.2), the model achieved an accuracy of 0.75 on the training dataset and 0.3 on the test dataset, as detailed in the confusion matrix (**Figure 5B**).

**Figure 5:**
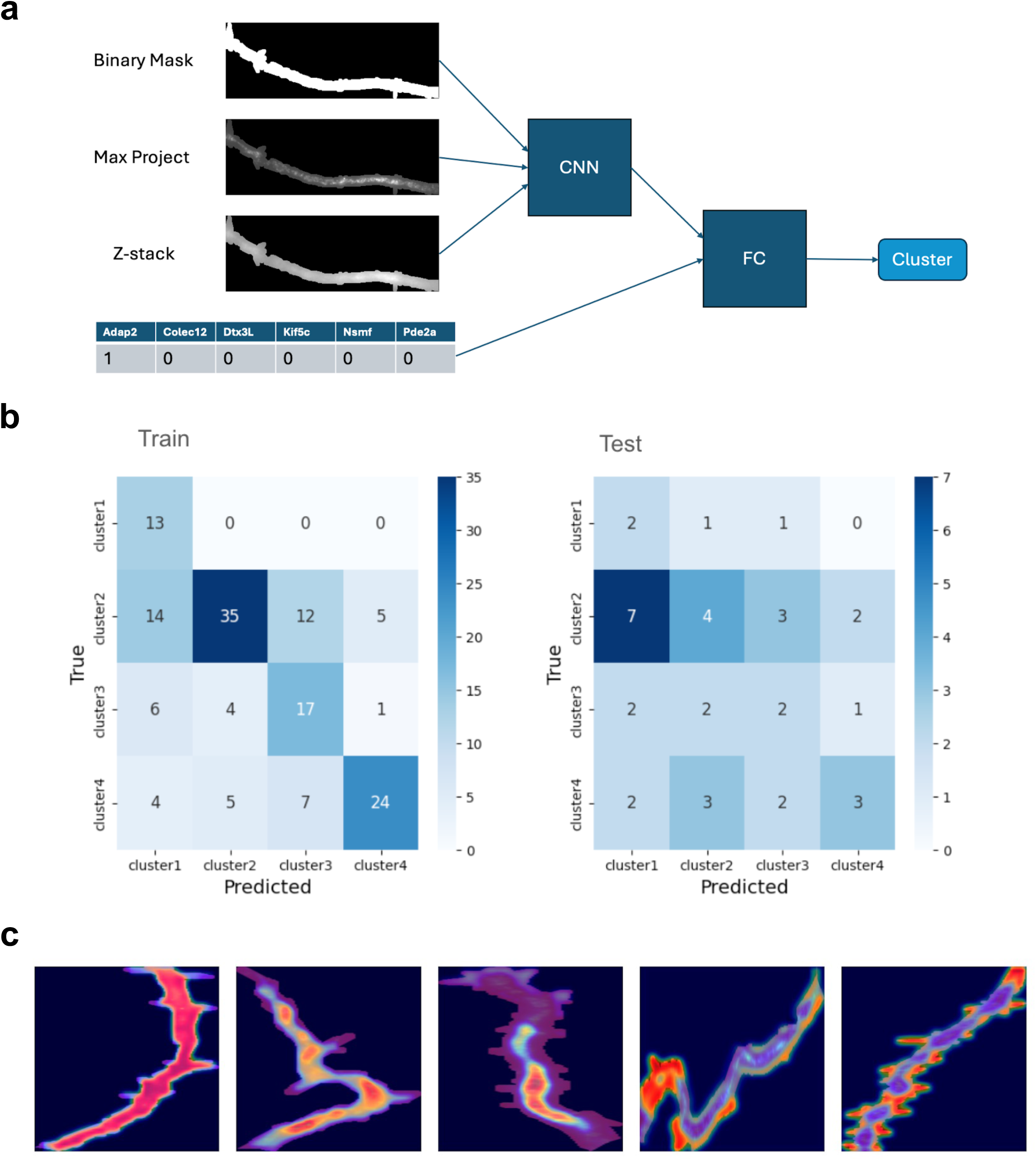
Analysis of dendritic morphology and mRNA spatial distributions using machine learning approaches. **A)** A schematic representation of the CNN model utilizing binary masks, Z-stacks, and maximum projection images of dendrites, integrated with one-hot encoded gene identity vectors, to predict cluster labels associated with mRNA distribution. **B)** Confusion matrices for the CNN classifier showing the performance on training and test datasets, highlighting the challenge of achieving high accuracy due to the unbalanced nature of the data. **C)** GradCAM visualizations demonstrating the convolutional layer activations associated with the various predicted clusters, which suggest patterns but do not conclusively identify specific morphological features across layers.

Although limited data restricted the development of a highly robust classifier, GradCAM analysis (Selvaraju et al. 2020) revealed various levels of pattern intensities within the convolutional layer activations, as shown in **Figure 5C**. This analysis highlighted differing scales, areas, and boundary details in the morphology that correspond to mRNA distribution labels. However, these morphological features did not consistently correlate with specific cluster labels across the dataset.

### Simulation of Passive Transport in 1D

Next, we modeled the passive transport of mRNA molecules within the dendrites and found that diffusion alone cannot reproduce the spatial patterns observed in the single dendrites. Currently, the known mechanisms of dendritic localization involve recognition of localization elements by transport proteins, formation into transport granules, and active transport along the microtubules by associated motor proteins (Mayford et al. 1996, Eom et al. 2003, An et al. 2008, Lee et al. 2016, Taliaferro et al. 2016, Tushev et al. 2018, Arora et al. 2022). However, localization of mRNA in other cell types relies primarily on passive diffusion (Forrest and Gavis 2003, Kugler and Lasko 2009, Yamagishi et al. 2009, Park et al. 2014, Engel et al. 2020). The genes selected for our study are known to be dendritically localized, but mechanisms underlying their localization are not well understood. Consequently, we assessed whether the observed spatial distribution of mRNA can be reproduced by a simple model of passive diffusion.

Briefly, we modeled the diffusion of mRNA molecules within dendrites as a one-dimensional random walk, as mRNAs were detected within 1 - 3 μm of the skeletonized center of dendrites (**Supplementary Figure 4**). Our model assumes both a constant diffusion coefficient across the length of the dendrite and constant rate of mRNA replacement. Previously reported diffusion coefficients of mRNAs vary widely, likely due to differences in measurement methods and genes probed (Fusco et al. 2003, Park et al. 2014, Fonkeu et al. 2019, Fujita et al. 2020); here, we used 1 μm^2^/second speed, which is in line with the most recently reported rate and corresponds to the upper bounds of the range of reported values. Previous simulations of diffusion in the dendrites (Fonkeu et al. 2019, Fujita et al. 2020) assume that *de novo* transcription is the primary source of mRNA in the dendrites, deriving the rate of transcription (0.001 mRNA per second) to represent the rate of introducing new mRNA molecules to the dendrite. By contrast, in our model, we assume the primary source of new mRNAs to the dendrites are pre-existing somatic mRNAs; as we observed a consistently higher density of mRNAs within the soma, we assume there is a constant replenishment. We simulated the random walks for each individual mRNA molecule for up to 30 minutes across 300 simulated dendrites, then compiled the simulated mRNAs’ distances from the soma at various time points. As the reported mRNA half-life range in the multi-hour scale (Schwanhäusser et al. 2011, Tushev et al. 2018), mRNA degradation was not considered in our model.

Generally, our compiled distribution of simulated mRNAs’ distances follows an exponential distribution with the highest mRNA density closest to the soma, as reported previously (Fonkeu et al. 2019, Fujita et al. 2020). The mRNAs’ furthest distance traveled increased over time, reaching ∼ 100 μm within 20 minutes, consistent with previous reports that newly transcribed mRNAs localize to the dendrites on the timescales of hours (Akbalik et al. 2017). The shape of the compiled distribution of simulated mRNAs’ distances did not align with the compiled distribution of observed mRNAs’ distances for any gene probed as the simulations had much sharper decay in mRNA density and could not reproduce the zone of low mRNA molecules immediately outside the soma (**Figure 6A**, **Supplementary Figure 3A**). Next, we computed the compiled distribution of simulated mRNAs’ *normalized* distances by dividing the simulated mRNAs’ distances from the soma by the length of the simulated dendrite. The resulting distribution from the simulations remains a clear exponential, with the decay rate of mRNA density decreasing as elapsed time increases. By contrast, the compiled distribution of normalized distances from the detected puncta, suggests that mRNAs are roughly uniformly distributed (**Figure 6B**, **Figure 4A**). Notably, in our simulations, the maximum mRNA density was found consistently immediately outside the soma in contrast to the observed spatial patterns from our single dendrites analysis, where the maximum mRNA density was at the center or the distal end of the dendrite for clusters **C1** and **C4** respectively. Taken together, our model demonstrates that diffusion alone is unable to reproduce the observed characteristics in the spatial patterning of mRNAs, likely indicating that our genes are organized by more active transport mechanisms.

**Figure 6:**
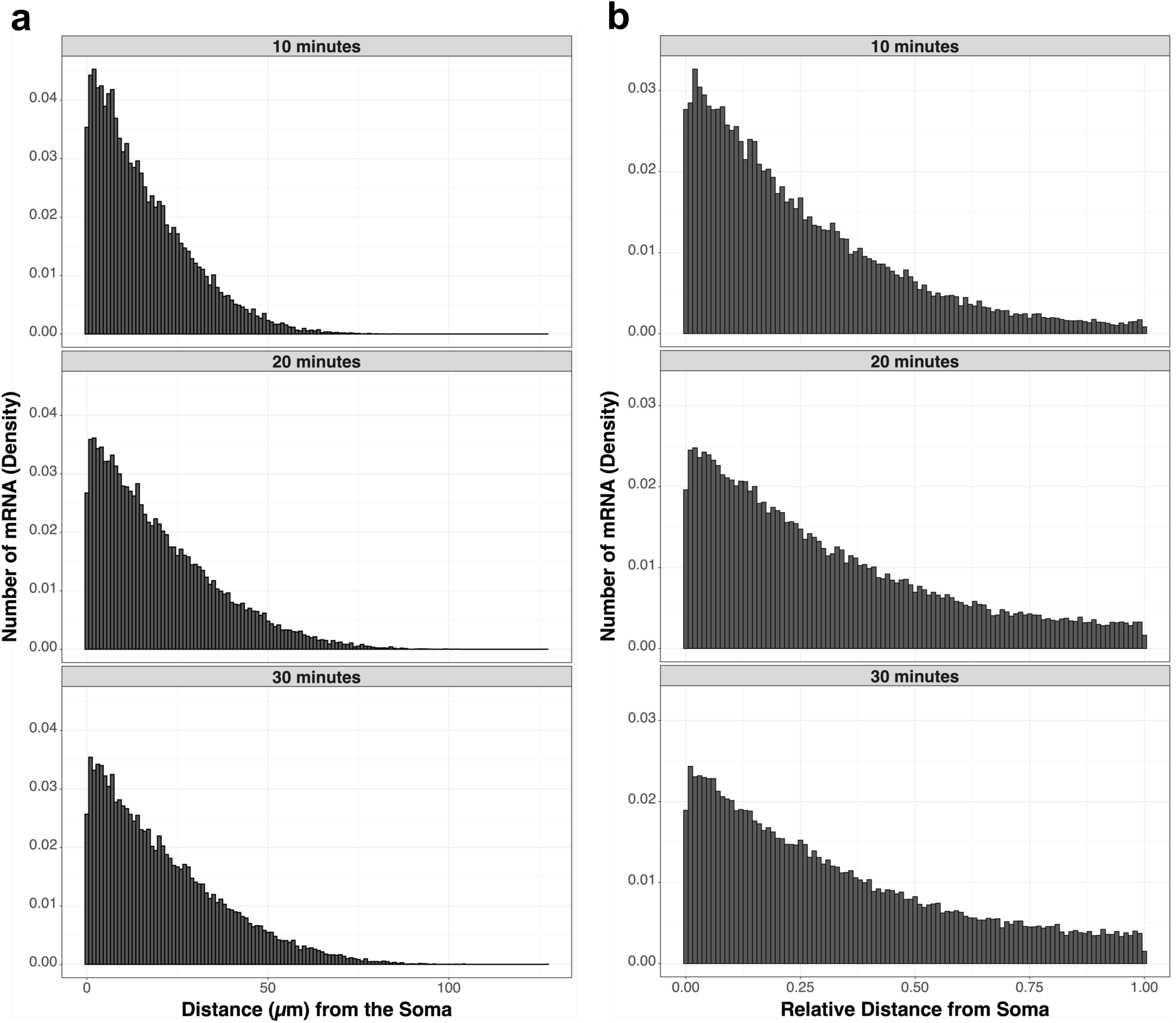
Simulation of passive transport in dendrites by random walk. Histograms of **A)** raw distances and **B)** *normalized* distances from the soma for simulated mRNAs. Each panel represents a snapshot at a specific time elapsed from the start of the simulation.

It should be noted that our model is not a comprehensive description of the RNA transport dynamics within the neurons; there are numerous active and passive factors that affect the observed spatial distribution in dendrites, such as local differences in viscosity, molecular compositions, and morphology (Sun et al. 2021), synaptic plasticity (Donlin-Asp et al. 2021), or RNA anchoring (Forrest and Gavis 2003), may affect diffusion rates or particle kinetics locally. However, our simplified model does recapitulate the spatial patterns of mRNAs predicted by more complex models: moreover, the effects of these factors would not result in distributions observed in passive transport.

### Colocalization of mRNAs

Colocalization of mRNA in dendrites has proven to be difficult to detect and define as the mechanisms of transport are still relatively unknown. One simple definition and model of colocalization is co-transport, which involves the assembly and transport within the same RNA carrying protein granules.

Previously, there have been conflicting reports, using the gene sets, of whether 1) mRNAs from different genes are co-transported and 2) multiple mRNA molecules of the same genes are co-transported within the same granule (Gao et al. 2008, Tübing et al. 2010, Mikl et al. 2011, Batish et al. 2012). Most recently, Batish et al. 2012 reported that there was little evidence of either: they used a threshold distance of 250 nm, which they estimated as the upper limit for the size of a granule, to compute the proportion of total mRNA molecules per gene that colocalized with another. Their observed percentages of colocalized mRNAs ranged between 0.33 - 3.36% for any combination of genes. However, when we repeated the analysis as specified in Batish et al. 2012, on our experimental data, we found that a much higher proportion of mRNA molecules were within the threshold distance, ranging from 2.58 - 23.02% across gene combinations (**Table 1**, **Supplementary Figure 5A**). These percentages are significantly different from those previously reported, suggesting there are higher incidences of colocalization events for the gene combinations in our study, despite similarities in experimental conditions. As Batish et al.’s analysis examines the distribution of nearest neighbor distances between mRNA molecules, we expect it to follow an exponential distribution, based on Extreme Value Theory, as observed in our distributions but not in Batish et al. (**Supplementary Figure 5B**). Furthermore, Mikl et. al 2011 previously described the proportion of colocalized molecules for any two dendritic mRNAs ranged from 5.7 - 8.3%, suggesting that the percentages reported in Batish et al. may be lower than expected.

**Table 1:** Colocalization frequencies of different gene combinations. Each cell presents the mean percentage of transcripts detected from transcripts from the row-labeled gene detected within 250 nm of transcripts from the column-labeled gene. Gene combinations of probes ordered in the same color channel could not be tested and left blank.

|  | *Adap2* | *Colec12* | *Dtx3L* | *Kif5c* | *Nsmf* | *Pde2a* |
| --- | --- | --- | --- | --- | --- | --- |
| *Adap2* | 20.23 ± 0.66 | 5.72 ± 0.4 |  | 7.33 ± 0.55 | 10.26 ± 0.51 | 3.73 ± 0.42 |
| *Colec12* | 16.53 ± 0.98 | 11.42 ± 0.62 | 11.42 ± 0.62 | 5.73 ± 0.72 | 8.31 ± 0.63 |  |
| *Dtx3L* |  | 3.97 ± 0.36 | 20.8 ± 0.62 | 7.54 ± 0.73 | 13.5 ± 1.44 | 2.83 ± 0.22 |
| *Kif5c* | 23.02 ± 1.2 | 5.83 ± 0.63 | 13.67 ± 0.77 | 23.68 ± 1.24 |  | 5.59 ± 0.86 |
| *Nsmf* | 19.56 ± 0.8 | 4.49 ± 0.36 | 15.9 ± 0.85 |  | 22.64 ± 0.7 | 2.58 ± 0.22 |
| *Pde2a* | 16.66 ± 1.06 |  | 12.95 ± 0.93 | 6.24 ± 0.58 | 9.43 ± 1.01 | 17.68 ± 1.01 |

To identify gene pairs that are likely colocalized, we focused on gene pairs with an observed total proportion of molecules colocalized above the 8.3% upper limit between any two dendritic RNA reported by Mikl et al. It is important to note that we measure the proportion of colocalized molecules in both directions, that is we computed both the proportion of mRNA molecules from gene A that were within 250 nm of gene B and the proportion of mRNA molecules from gene B that were within 250 nm of gene A. From our analysis, *Adap2* and *Nsmf* and *Dtx3L* and *Nsmf* were both reciprocally found within 250 nm of each other 10 - 19%, while for other combinations (*Colec12* and *Dtx3L*; *Colec12* and *Adap2*; *Kif5c* and *Adap2*; *Kif5c* and *Dtx3L*; *Pde2a* and *Adap2*; *Pde2a* and *Dtx3L*; *Pde2a* and *Nsmf*), same range of high proportion of colocalization was only found in one direction. Generally, the proportion of colocalized molecules was not reciprocal across gene pairs, which is to be expected given the differences in gene expression levels. This suggests that *Adap2* and *Nsmf* and *Dtx3L* and *Nsmf* may be co-transported within the same granule. As probes for *Adap2* and *Dtx3L* were in the same color channel, we could not assess whether they would also colocalize.

*Adap2*, *Dtx3L*, and *Nsmf* were each from different functional classes miscellaneous, immune response, and neuronal respectively, suggesting gene functionality is not a good predictor of colocalization. However, given our limited gene set, a more comprehensive work using genes from well-established gene networks will be required to better address whether genes within the same functional class are likely to be colocalized.

From our analysis, the observed proportions of molecules within 250 nm, we note that, except for *Colec12*, there is a higher proportion of colocalization events for mRNA molecules from the same gene, ranging from 17.68 - 23.68% (**Table 1**), compared to proportion of colocalization events of mRNAs from different genes, which suggests that multiple mRNA molecules of the same gene may be co-transported. This corroborates the finding in Miki et. al. which also reported higher incidence of colocalization proportions for mRNA molecules from the same genes, ranging from 10.5 - 28%. This likely indicates that one to two mRNA molecules from the same gene are co-transported for all genes in our study, though as our observed percentages are generally larger than those reported in Mikl et al. study, the likelihood of a granule having two copies may be higher.

Co-expression and concurrent expression of genes required for remodeling of synaptic spines and sub-dendritic arbors can range from 0.5 μm to 5 μm (Sun et al. 2021). Therefore, there is a need to characterize colocalization events across a wider range of distances than the size of an RNA granule. To address colocalization patterns at large spatial scale, we calculated Ripley’s K of the mRNAs’ physical locations within single dendrites, which involves calculating the pairwise distances between mRNA molecules of the same gene (self-colocalization) and across gene combinations (multi-gene colocalization) to construct the cumulative distribution of colocalization events at search radii ranging from 0.1 μm to 6 μm (**Figure 1**). To assess the significance of the colocalization frequencies, we calculated the Ripley’s K of 1000 simulations of uniformly distributed points, equal to the number of mRNAs to the analyzed dendrite, then used the 5^th^ and 95^th^ percentile values to construct a confidence envelope at each interaction radius (**Figure 1**, **Methods**). To accurately characterize colocalization, we only calculated the Ripley’s K of dendrite-gene pairs with at least 30 detected mRNA molecules of a gene when analyzing for self-colocalization and dendrite-gene combination pairs with at least 20 detected mRNA molecules of each gene when analyzing multi-gene colocalization. At each interaction radius, we tallied the number of dendrite-gene or dendrite-gene combination pairs, where the observed Ripley’s K was larger than the 95^th^ percentile then subtracted the number of pairs where the observed Ripley’s K was lower than the 5^th^ percentile.

Generally, we find that the colocalization patterns were comparable across all gene combinations, including self-colocalization. We found that colocalization occurred frequently within 0 - 2 μm, beyond what was expected from our simulation, consistent with both our previous colocalization analysis using nearest neighbors and previous reports of colocalization by co-transport (**Figure 7A**, **Figure 7B**). This supports our previous conclusion that multiple genes may be co-transported and that multiple copies of the same gene can be packaged into the same granule. Additionally, as the estimated upper limit of the size of transport granules in neurons is 600 - 700 nm (Barbarese et al. 1995), our results suggest that the transport of multiple different granules could be regulated so that each granule ends up in close proximity to one another.

**Figure 7:**
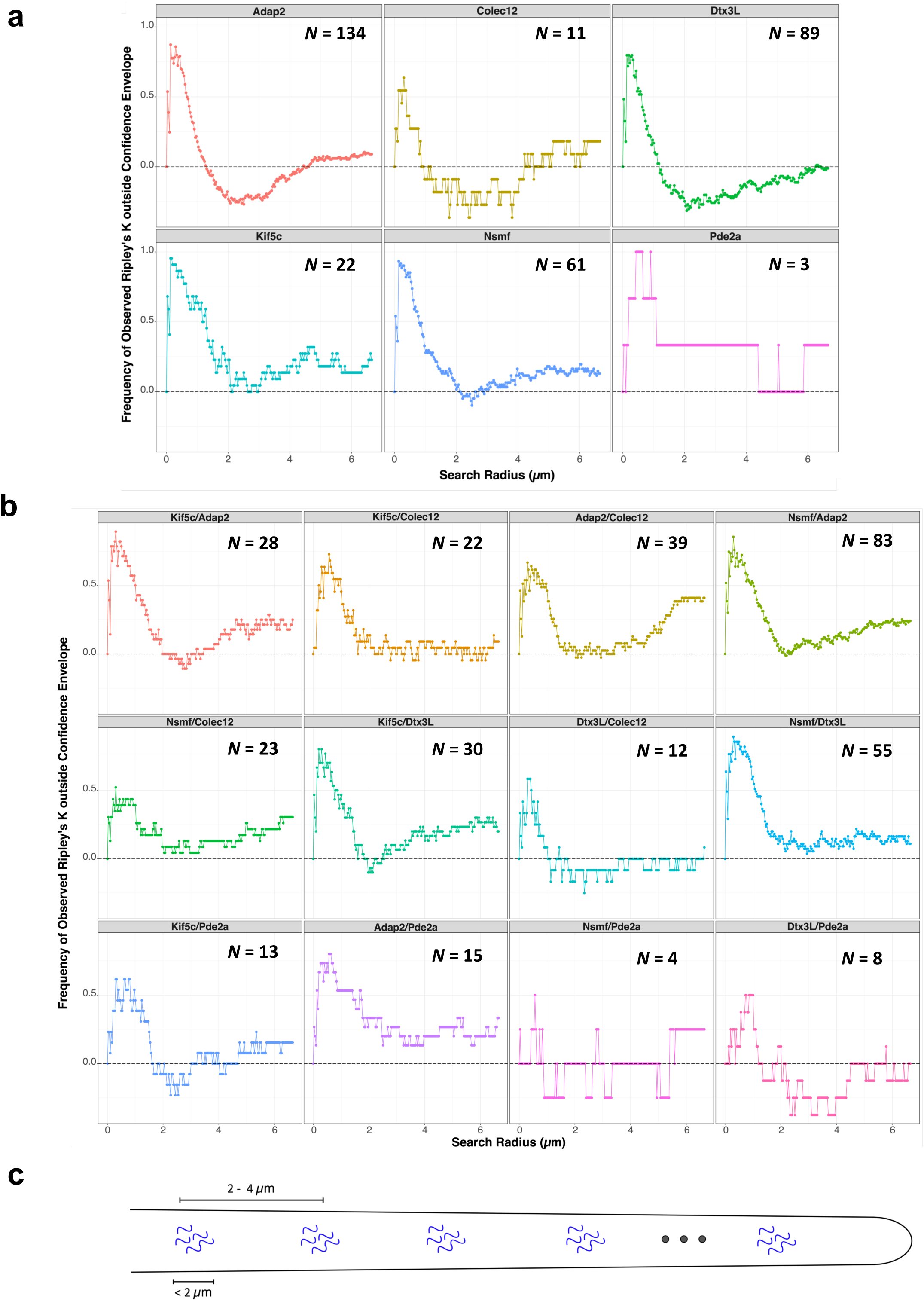
Characterizing colocalization frequencies of dendritically localized mRNAs at various search radii. Scatter plots present summarized Ripley’s K (number of samples with observed Ripley’s K larger than the 95^th^ percentile subtracted by the number of samples with observed Ripley’s K was lower than the 5^th^ percentile) at each radius. Each panel represents **A)** genes in analyses of self-colocalization or **B)** gene combinations in analyses of multi-gene colocalization. *N* represents the number of dendrite-gene/dendrite-gene combination pairs used in each analysis. **C)** Diagram of spatial organization of dendritically localized mRNAs derived from Ripley’s K analysis.

Notably, frequency of colocalization events peaks at and returns to ranges expected from our simulation at different interaction radii across gene combinations, possibly hinting at differences in the sizes and number of the transport granules across genes or in the size of the affected downstream processes. Next, in the range of interaction radii 2 - 4 μm, we found that frequencies of self-colocalization events for *Adap2*, *Colec12*, and *Dtx3L* were low, below the expected range of our simulations, suggesting that the clusters of mRNA molecules are overly dispersed at these distance scales (**Figure 7A**). A similar tendency was observed for gene combination *Kif5c*/*Pde2a* in our multi-gene colocalization analysis, though not as strongly (**Figure 7B**). The remaining gene combinations, including self-colocalization, were within expectations of our simulations. Lastly, in the range of interaction radii 4 - 6 μm, we found that the frequencies of colocalizations for gene combinations *Kif5c*/*Adap2*, *Adap2*/*Colec12*, *Nsmf*/*Colec12,* and *Kif5c*/*Dtx3L* were higher than the expected range; these colocalization frequencies at interaction radii from 4 - 6 μm were about ⅓ - ½ of the colocalization frequencies at interaction radii from 0 - 2 μm (**Figure 7B**) A similar tendency was observed in self-colocalization for genes *Adap2*, *Kif5c*, and *Nsmf* (**Figure 7A**). The remaining genes and gene combinations were within expectations of our simulations.

Our findings suggest that neuronal mRNAs within individual dendrites are organized into colocalized clusters at 0 - 2 μm scale, then randomly spaced at higher distance scale; but for certain genes, there was evidence of over-dispersion (i.e., evenly spaced) at 2 - 4 μm scales (**Figure 7C**). These patterns themselves may be grouped into spatial patterns that are larger scaled. Taken together, our analysis suggests evidence of colocalization beyond the upper limits of RNA granule size. Our analyses of colocalization suggest regulated multi-scale spatial structure of mRNAs within the dendrites, which support previous reports of subdomains within the dendrites (Sun et al. 2021) and motivates further studies of larger spatial organization of mRNAs within dendrites.

## DISCUSSION

RNA localization in dendrites is key to providing the neuron with timely local translation of protein for synaptic modulation, and its disruption has been linked to neurodegeneration (Sutton and Schuman 2006, Buxbaum et al. 2015, Holt et al. 2019, van Spronsen and Hoogenraad 2010, Penzes et al. 2011, Swanger and Bassell 2013). The study of fine spatial distributions of individual mRNA species within the dendrites is important to elucidating the roles of the extensive catalog of dendrite localized or enriched genes already identified by previous work (Zhong et al. 2006, Poon et al. 2006, Ainsley et al. 2014, Taliaferro et al. 2016, Middleton et al. 2019). In this study, we systematically analyzed the spatial distribution of dendritic mRNAs from different genes by multiplexed single molecule FISH. This approach is more robust at detecting mRNA than sequencing workflows, which is prone to both failure at capturing low-abundance RNA molecules and bias in counts from PCR amplification (Torre et al. 2018), and more importantly, provides greater spatial resolution. The complex morphology of dendrites poses a significant challenge for characterizing accurate position of individual mRNA molecules, which previous methods have addressed by linearizing the dendrites in their image analysis (Cajigas et al. 2012, Buxbaum et al. 2014). Here, we employ a graph-based analysis to navigate the geometry of dendrite and characterize the density of s detected with smFISH as a function of the distance from the soma.

Our results suggest that both dendritic morphology and gene identity contribute to determining the spatial distribution of mRNAs within the dendrites. Previous studies have characterized the spatial distribution of the mRNA molecules from specific genes along the dendrites (Cajigas et al.2012, Buxbaum et al. 2014, Chen et. al 2020), typically binning the number of mRNA molecules at 10 - 25 μm intervals. Our graph-based methodology allows for measurements of distances up to pixel level resolution, around 133 nm, improving the precision of our spatial measurements.

Our results support the previous findings (Cajigas et al. 2012) that, largely, distributions of mRNAs within the dendrites vary in the initial concentration at the proximal dendrite and the rates of decay of mRNA density over the length of the dendrites. However, as evident in the single-dendrite analysis, we found that spatial distribution of mRNAs was sensitive to the morphology of the dendrites, specifically the length. Our analysis demonstrated that mRNA molecules were generally present throughout the entire length of the dendrite and the location of the maximum density of mRNA is not limited to the proximal end of the dendrite.

We found that genes were not limited to a single spatial pattern, though some genes are found more frequently within certain patterns. Our analysis suggests there is an interplay between dendrite morphology and gene identity that affects the mRNAs’ spatial distribution within the dendrite. Gene function and activation states of the dendrites may further influence dendritic localization dynamics.

Given our limited dataset, our Machine Learning models did not well-characterize common features that might determine patterns of spatial localization. A more thorough investigation of dendritic morphology and the spatial pattern of localized genes may reveal functionally relevant morphological features that affect dendritic localization.

While our study does not directly elucidate additional mechanisms of RNA localization, we found that a model of passive diffusion was insufficient to explain two observed characteristics of the mRNAs’ spatial pattern: the zone of low mRNA density immediately outside the soma and 2) variability in the location of the highest mRNA density. Our results highlight the importance of active transport mechanisms in regulating the spatial distribution of dendritically localized mRNAs.

As discussed previously, there have been conflicting reports of whether mRNA molecules from different genes or multiple mRNA molecules from the same gene are packaged within the same granule for co-transport (Gao et al. 2008, Tübing et al. 2010, Mikl et al. 2011, Batish et al. 2012). In our study, we found evidence of colocalization for at least 2 gene pairs, Adap*2-Nsmf* gene pair and *Dtx3L*-*Nsmf* gene pair, which were reciprocally found within 250 nm of each other, possibly indicating co-transport within the same granule. Furthermore, we observed a higher proportion of colocalization for mRNA molecules from the same gene, pointing to higher incidence of granules with two mRNA molecules from the same gene. While our results do not address the ambiguities in the genes previously studied, as different groups of genes likely demonstrate different patterns of colocalization, it does present evidence for co-transport both within and across genes.

We characterized the statistical significance of colocalization frequencies between mRNAs at various interaction radii with Ripley’s K and found that neuronal mRNAs within individual dendrite are organized into clusters at short length scale, which in turn are distributed uniformly randomly or in some cases evenly. It still remains unclear if colocalized mRNAs at scales beyond co-transport are meaningful to the dendritic spatial organization of dendrites, though regulation of localization mechanisms could lead to many different granules being colocalized without co-transport. Additionally, larger distance colocalization may reflect subtle morphological features within the dendrites such as synaptic neighborhoods (Sun et al. 2021), which may be centralized loci for RNA capture or correlates with the area of the downstream processes affected by the co-expression of the colocalized mRNAs.

Although we examined a limited number of genes, we introduce analysis tools to explore how mRNA spatial patterns correlate with the key morphological features of dendrites. Our study highlights the effects of morphology on the spatial distribution of dendritically localized mRNAs. It is still unknown how specific morphological features interact with transport mechanisms to affect localization, nor the role of these morphological features in the larger cellular functions of the neuron. Additionally, as a recent study (Chen et al. 2021) reported that different isoforms of the same genes displayed different spatial distributions within the dendrites, a more systematic assay with higher degree of multiplexing with single molecular resolution will be required to fully dissect the density distributions associated with each isoform.

## METHODS

### smFISH probe design

We designed custom Stellaris FISH probes using the Stellaris RNA FISH probe designer (LGC Biosearch Technologies) and mRNA sequences from MGI (Blake et al. 2021) with care to avoid targeting mRNA splice junctions. Probe sets with reporter dyes Cal Fluor Red 610 (*Adap2*, 48 probes; *Dtx3L*, 48 probes), Quasar 570 (*Colec12*, 48 probes; *Pde2a*, 48 probes), and Quasar 670 (*Kif5c*, 48 probes; *Nsmf*, 48 probes) were purchased from Biosearch Technologies.

### Neuron culture

Hippocampal neurons from embryonic day 18 (E18) mice (C57BL/6) were cultured for 15 days, as previously described (Buchhalter and Dichter 1991). Briefly, neurons from dissociated hippocampus were plated onto acid washed, poly-D-lysine coated 12 mm growth coverslips (Neuvitro) at concentrations between 80,000 - 100,00 cells/mL. Cultures were grown in Neurobasal medium (Gibco) supplemented with 2% B-27 and 0.26% Glutamax (200 mML-alanyl-L-glutamine dipeptide in 0.85% NaCl) at 37°C and 5% CO_2_. Experiments were performed at 15 days in vitro (DIV).

### Multiplexed single molecule RNA fluorescence in situ hybridization (smFISH)

At 15 DIV, neurons were fixed in 4% paraformaldehyde in RNAse-free phosphate buffered saline (PBS) for 10 minutes. Fixative was removed with serial PBS washes. Cells were then permeabilized and stored in 70% ethanol at 4°C overnight. The following day, coverslips were rehydrated by serial PBS washes at room temperature, then washed in 2X saline-sodium citrate (SSC) buffer to enhance RNA detection for no more than 30 minutes. Each coverslip was flipped onto a drop of 50 µl of hybridization solution on parafilm; hybridization solution per 12 mm coverslip contained: 1.5 μl of Stellaris smFISH probes (stock molarity of 12.5 μM), 1 μl of primary antibody targeting MAP2 (concentration: 1 μg/μL, Abcam Cat.# ab32454) in 45 μL of hybridization buffer (BioSearch Cat.# SMF-HB1-10) and 5 μL of formamide (Fisher Cat.# BP227-500). Hybridization proceeded overnight at 37°C in a sealed humidified chamber. The following day, coverslips were transferred into a 12-well dish and washed in 1 mL of wash buffer (5 mL formamide, 5 mL 20X SSC, 40 mL DEPC-treated H_2_O) containing 1 μL of secondary antibody (concentration: 2 μg/μL, Abcam Cat.# ab150077) for 30 minutes, twice at 37°C. DAPI for nuclear staining was added in the secondary wash. Coverslips were mounted on PBS for imaging.

### Fluorescent Imaging of smFISH stained neurons

Images were taken on Zeiss Axio Imager M2 microscope equipped with an X-Cite Series 120PC Q lamp, using 100X PlanApoChromat 1.40 NA oil immersion objective (Zeiss) and Andor iXon 888 EMCCD camera. 400 nm step Z-stack images (15 planes) were taken with the following exposure times: 3 - 5 ms (DAPI), 150 - 250 ms (GFP) and 2,000 - 3,000 ms (Cy3, Cy5, mCherry) using MetaMorph software (Molecular Devices) at 0.133 µm/pixel (**Figure 1**). Care was taken to image neurons that were clearly distinguishable by cell body height and process morphology, with healthy distinct dendrites without fasciculation. A total of 410 neurons were imaged from multiple cultures across eight triplet probe combinations on multiple days.

### Morphological characterization and puncta detection

We used SynD (Schmitz et al. 2011) to analyze the GFP stained neuron to detect and separate the soma and dendrites (**Figure 1**). The SynD generated masks were binarized in ImageJ (Schneider et al. 2012). To ensure that the soma masks are properly connected with the dendritic masks and exclude regions of ambiguity at the base of the dendritic masks, we dilated the somatic masks using Python OpenCV (Bradski 2000). Additionally, we removed any small regions of dendritic masks that did not extend outside the expanded somatic masks and excluded them from any future analysis. The location of each single-molecule FISH puncta was identified using Aro (Wu and Rifkin 2015); briefly, potential puncta were initially identified by locating local intensity maxima, then processed through a user-trained, supervised random forest classifier (**Figure 1**). Training image sets for the random forest classifier were created specifically for each triplet probe combination to account for variability in staining. The pixel coordinates of each detected puncta were extracted with a custom MATLAB code, which was used to classify the puncta as somatic or dendritic based on whether the puncta coordinates were within the soma or dendrite binary mask.

### Graph-based distance calculation

To best calculate the distances between mRNA molecules within the complex geometry of the neuron morphology, we employed graph-based analysis using the networkX package (Hagberg et al. 2008) in Python (**Figure 1**). For each neuron, we created three classes of nodes: 1) *transcript nodes*, which represent the detected mRNAs within the dendrites, 2) *morphology nodes*, which represent the linear, skeletonized morphology of the dendrites, and 3) *edge nodes*, which represent the perimeter of the soma. Transcript nodes were assigned the pixel coordinates of their representative mRNAs. The dendrites mask was skeletonized using the scikit-image package (van der Walt et al. 2014) in Python. 50% of the resulting points were randomly selected, and the selected pixel coordinates were assigned as morphology nodes. The edge of the soma mask was detected by image convolution in the SciPy package (Virtanen et al. 2020); 25% of the detected points were randomly selected, and the selected pixel coordinates were assigned as edge nodes. All the nodes in the neuron were connected to their respective *k*-nearest neighbors, using Euclidean distances between their pixel coordinates, incrementing *k* until a single connected graph was constructed. For each transcript node, the distance to the soma was calculated as the minimum graph distance between the transcript node and the closest edge node by Dijkstra’s Algorithm (**Figure 1**), then converted to physical distances using the physical dimensions of the pixel (0.133 µm/pixel). The distances from the soma were partitioned into bins of 1 µm for the histogram plots (**Figure 1**).

### Single dendrite normalized distance analysis

To generate binary masks of single dendrites, we visually inspected every binarized dendrite mask and selected coordinates corresponding to well-separated single dendrites with minimal projections, no branching, and no overlap with other dendrites using the LassoSelector from the matplotlib Python package (Hunter 2007).

Distances from the soma were calculated as described above with one modification: the furthest morphology node from the perimeter of the soma (represented by edge nodes) was designated as the *end node*. The length of the single dendrite was calculated as the minimum graph distance between the end node and the edge nodes by Dijkstra’s Algorithm and used to normalize the raw distance from the soma into relative position along the dendrite. The normalized distances from the soma were partitioned into bins of 0.01 µm for the histogram plots.

### GradCAM

Grad-CAM, short for Gradient-weighted Class Activation Mapping, is a visualization technique for deep learning models (Selvaraju et al. 2020). It operates by producing “heatmaps” of class activation over input images, offering insights into which parts of the image are significant for making specific classification decisions. This technique employs the gradients of any target class flowing into convolutional layers to produce a localization map, highlighting the important regions for predicting the class.

In our study, we employed Grad-CAM to understand and validate the decision-making process of our CNN model to see which type of patterns emerged during the training. The core idea is to represent the spatial locations in convolutional feature maps by their importance to the particular class. These heatmaps are then overlaid on the input images, thus providing a visual explanation of the model’s decision-making process.

### Diffusion Simulation

To simulate the diffusion of mRNA molecules within dendrites as 1-dimensional random walk, we created two object classes in Python: a *transcript* class and a *dendrite* class. The transcript class had the following attributes: 1) position, which represents a mRNA molecule’s current position, 2) maximum distance, which represents the distance past which the mRNA cannot travel, and 3) half-life, which represents the time remaining before the mRNA is considered degraded and removed from the simulation. The dendrite class had the following attributes: 1) length, which represents dendrite length, 2) transcript array, a one-dimensional array of transcript objects, which represents the set of mRNA molecules within the dendrite. Each instance of a dendrite object was assigned a dendrite length and number of mRNAs, which were drawn randomly from the set of observed single dendrite lengths and number of mRNAs from our single dendrite analysis upon instantiation; the corresponding number of transcript objects were instantiated and added to the dendrite object’s transcript array. Upon instantiation, each instance of a transcript object had its position set to 0 and its maximum length set to the length of the dendrite it belongs to. To simulate random walk, each transcript object has an equal probability of moving to the left or right at every iteration. At the boundaries of the dendrite, the transcript object has an equal probability of staying or moving away from the boundary. The step-size moved per iteration was calculated by taking the square root of the diffusion coefficient (1 μm^2^/second) and multiplying by the time step (0.005 seconds), the amount of time represented by each iteration. At each iteration, every transcript object in the dendrite object’s transcript array underwent a random walk. As live dendrite would still be receiving mRNAs from the soma, both as novel transcription and existing somatic mRNA, new transcript objects were instantiated and added to the dendrite object’s transcript array at the mRNA introduction rate. As estimating the mRNA introduction rate is difficult, given its likely dependence on multiple factors, we tested a range of values from 0.05 – 0.1 mRNA per second; these values primarily affected the rate of decay in RNA density and the mean transcript distance from the soma and did not alter the overall conclusion that the simulations could not reproduce the observed characteristics in the spatial patterning of mRNAs. For **Figure 6**, we used the mRNA introduction rate of 0.083 mRNA per second, which closely matched the compiled distribution of mRNA distances for *Pde2a* (**Figure 3A**). Simulation continued until the final iteration number, calculated by the ending time point in seconds divided by the time step, was reached. A total of 300 dendrites were simulated for this analysis.

### Ripley’s K analysis for colocalization

For each single dendrite, transcript nodes, edge nodes, and morphology nodes were created and connected to their respective *k*-nearest neighbors to construct a single connected graph, as described above. Pairwise graph distances were calculated between all detected mRNAs, for analysis of both self-colocalization and multi-gene colocalization, using Djikstra’s algorithm (**Figure 1**). The frequencies of colocalization events at different length scales were calculated as the number of pairwise graph distances below various search radii (0 to 6 µm), and the cumulative distribution was plotted as the observed Ripley’s K (**Figure 1**).

To generate the confidence envelope, we generated the Ripley’s K of 1000 simulations of uniformly distributed points, representing mRNAs, and took the 5^th^ and 95^th^ percentile colocalization frequencies, for the lower and upper bounds respectively, at each search radius. For each simulation, we generated a one-dimensional line of equal length to and containing an equal number of points (mRNAs) as the analyzed dendrite. The coordinates of each point were assigned as follows: the x coordinates were drawn uniformly from the length of the dendrite and the y coordinates were drawn from a distribution of deviations from the center of the skeletonized line of the dendritic morphology (**Supplementary Figure 4A**). Pairwise Euclidean distances were computed across the simulated points and used to calculate the frequencies of colocalization at various search radii, as described above.

## DATA AVAILABILITY

Image data is available for download upon request.

## ACKNOWLEDGEMENTS

We thank Dr. Gregory Grant (University of Pennsylvania) for advice on statistical analysis.

