## Supplementary Figures for "Characterizing the Spatial Distribution of Dendritic RNA at Single Molecule Resolution"

#### SUPPLEMENTARY FIGURE LEGENDS:

**Supplementary Figure 1:** A comparison of the number detected mRNA molecules across probe combinations and neuronal compartments **A)** A violin plot of number of detected mRNAs across the triplet probe combinations. Probe combinations are shown in different colors. **B)** A violin plot of neuronal compartment area in pixels across the triplet probe combinations. **C)** A scatter plot of neuronal compartment area vs. detected mRNA molecules.

**Supplementary Figure 2:** Characterizing common spatial patterns of dendritically localized mRNAs. **A)** A quantile-quantile plot comparing the distributions of mRNAs' distances from the soma across genes, indicated by the row and column label. Each panel represents a pairwise comparison. **B)** Heatmap of pairwise Jensen-Shannon distances, which quantify the similarity, between the distance distributions of each neuron-gene pair. Hierarchical clustering of these distances is depicted in the dendrogram at the top of the heatmap. Each spatial patterning cluster identified is colored separately. **C)** Violin plot of dendrite compartment area of neurons assigned to each spatial patterning cluster identified. **D)** Bar graph representing the proportion of distance distributions from each probe combination that was assigned to each spatial patterning cluster. Probe combinations are represented in separate colors.

**Supplementary Figure 3:** Characterizing common spatial patterns of mRNAs within single dendrites. **A)** Histograms of graph distances from the soma of localized mRNAs within single dendrites. Binwidth is 1 µm. **B)** A quantile-quantile plot comparing the distributions of mRNAs' distances from the soma across neuron-level and single dendrite-level analyses. **C)** Heatmap of pairwise Jensen-Shannon distances, which quantify the similarity, between the distance distributions of each neuron-dendrite-gene combination. Hierarchical clustering of these distances is depicted in the dendrogram at the top of the heatmap. Each spatial patterning cluster identified is colored separately. **D)** Scatter plot of localized mRNAs' distances from the soma per each spatial patterning cluster. Each row of the scatter plot represents a single neuron-dendrite-gene combination. **E)** Violin plot of the number of mRNA molecules detected in the neuron-dendrite-gene combinations assigned to each spatial patterning cluster identified. **F)** Bar graph representing the proportion of distance distributions from each probe combination that was assigned to each spatial patterning cluster. Probe combinations are represented in separate colors.

**Supplementary Figure 4:** A histogram of mRNA molecules' distance to the center of the skeletonized center of the dendrites.

**Supplementary Figure 5:** Characterizing the colocalization frequencies of dendritically localized mRNAs. **A)** Each panel presents the jitter plot displaying the percentage of mRNA molecules from the gene indicated by the row label that were detected within 250 nm of mRNA molecules from the gene denoted by the column label. Probe combinations are represented by different colors. Mean percentages are summarized in **Table 1**. **B)** Each panel presents the histogram of nearest neighbor distances between mRNA molecules from the gene indicated by the row label to mRNA molecules from the gene denoted by the column label.

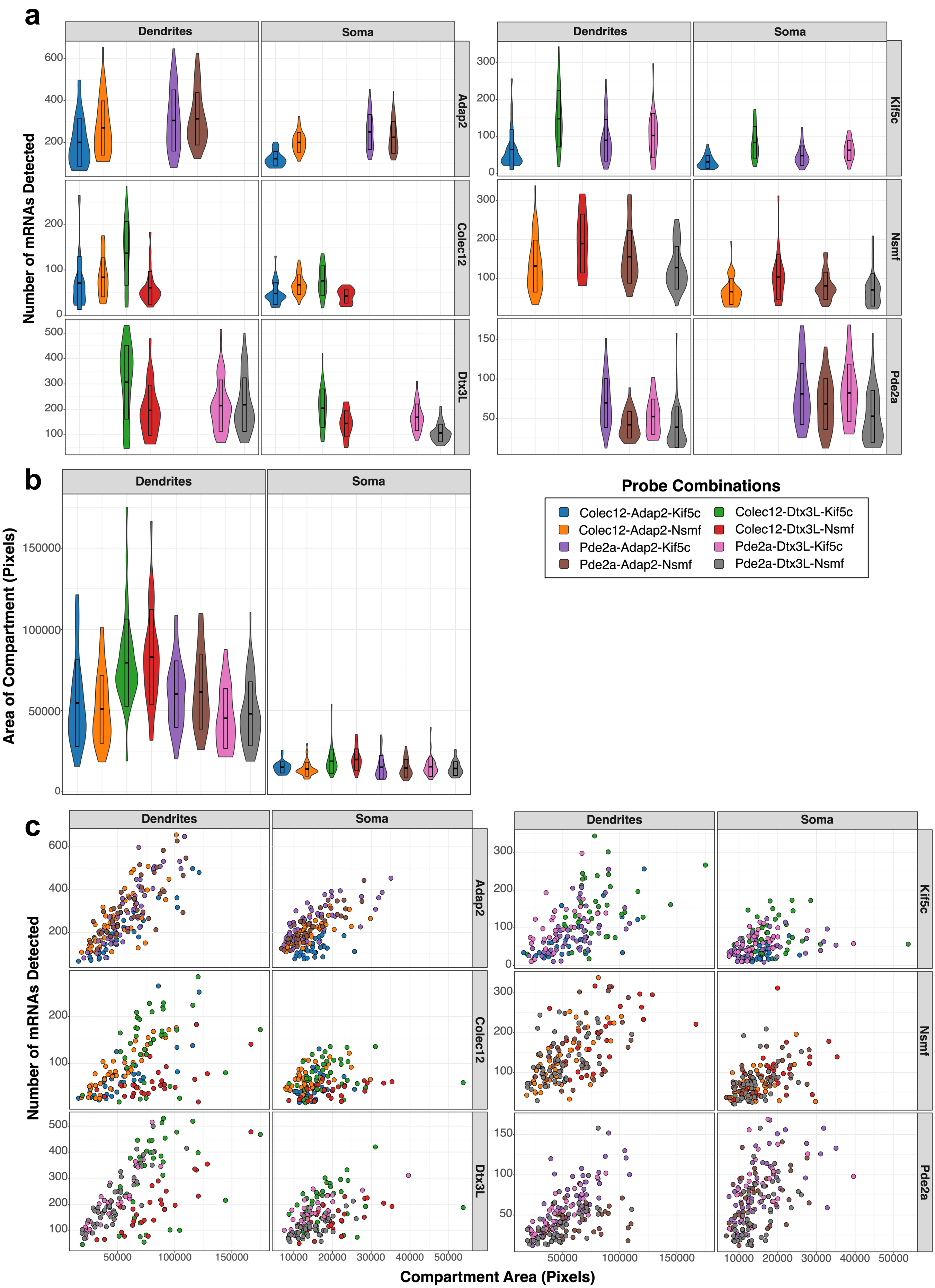


Supplementary Fig. 1

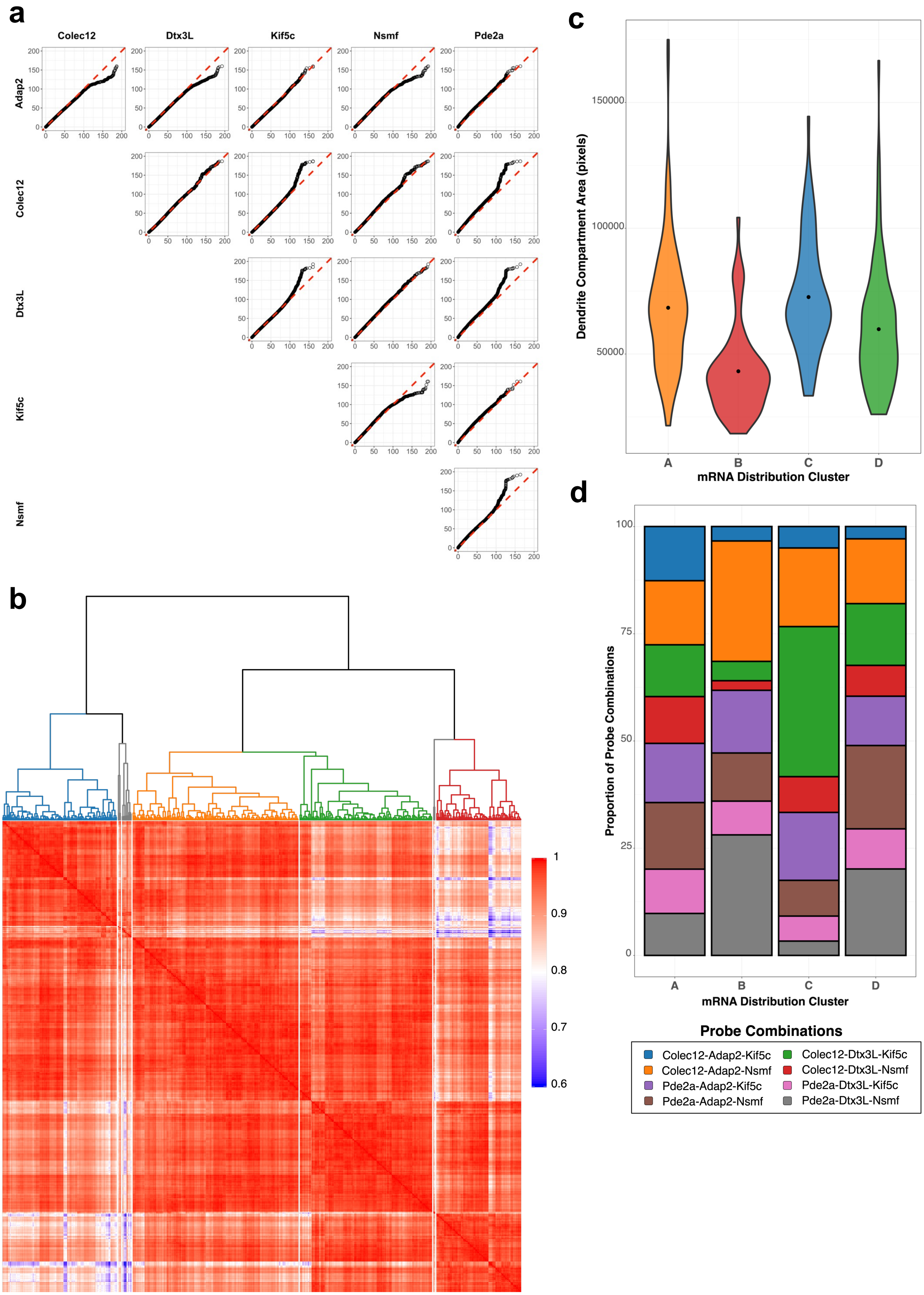


**Supplementary Fig. 2**

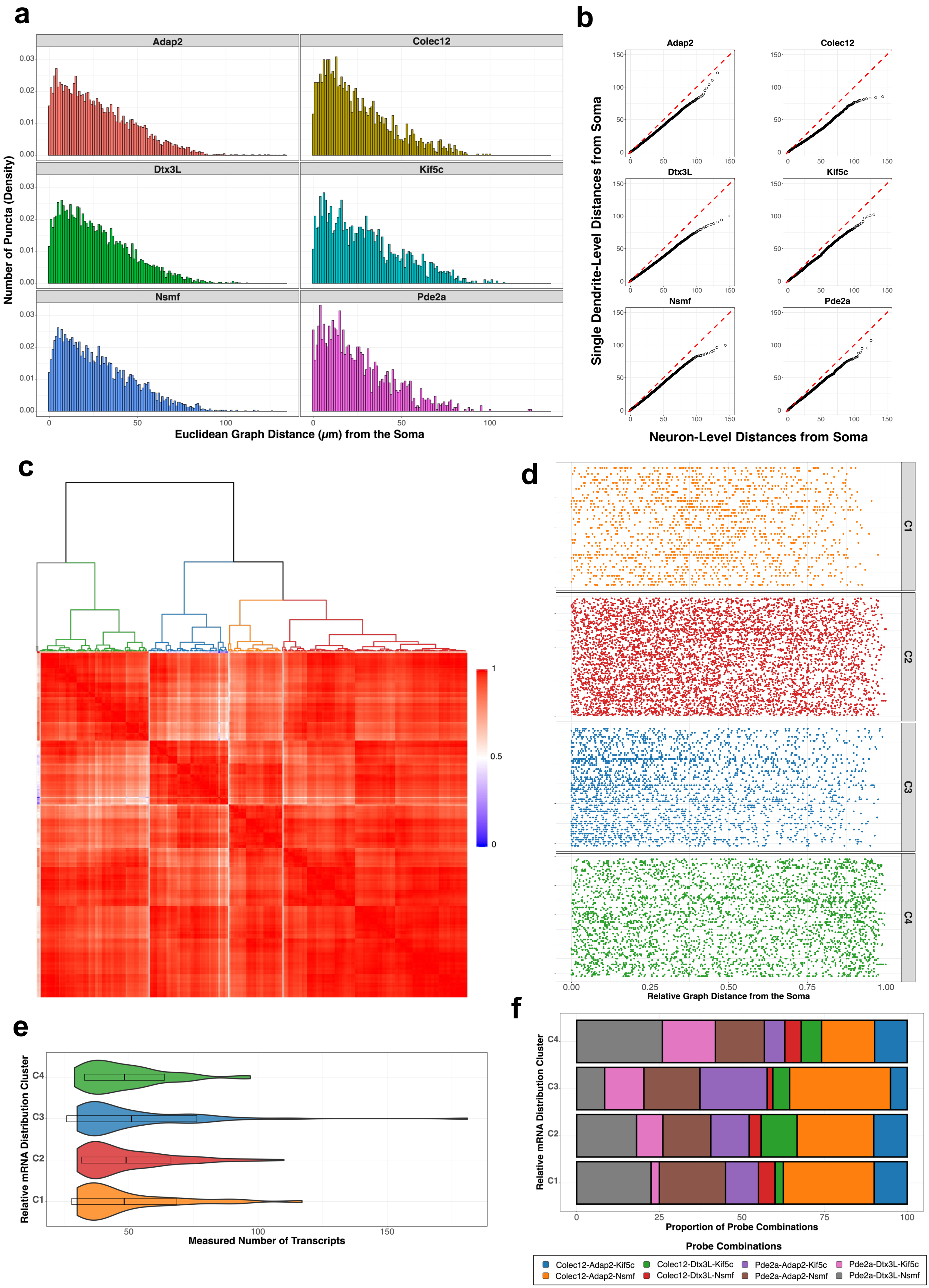


### Supplementary Fig. 3

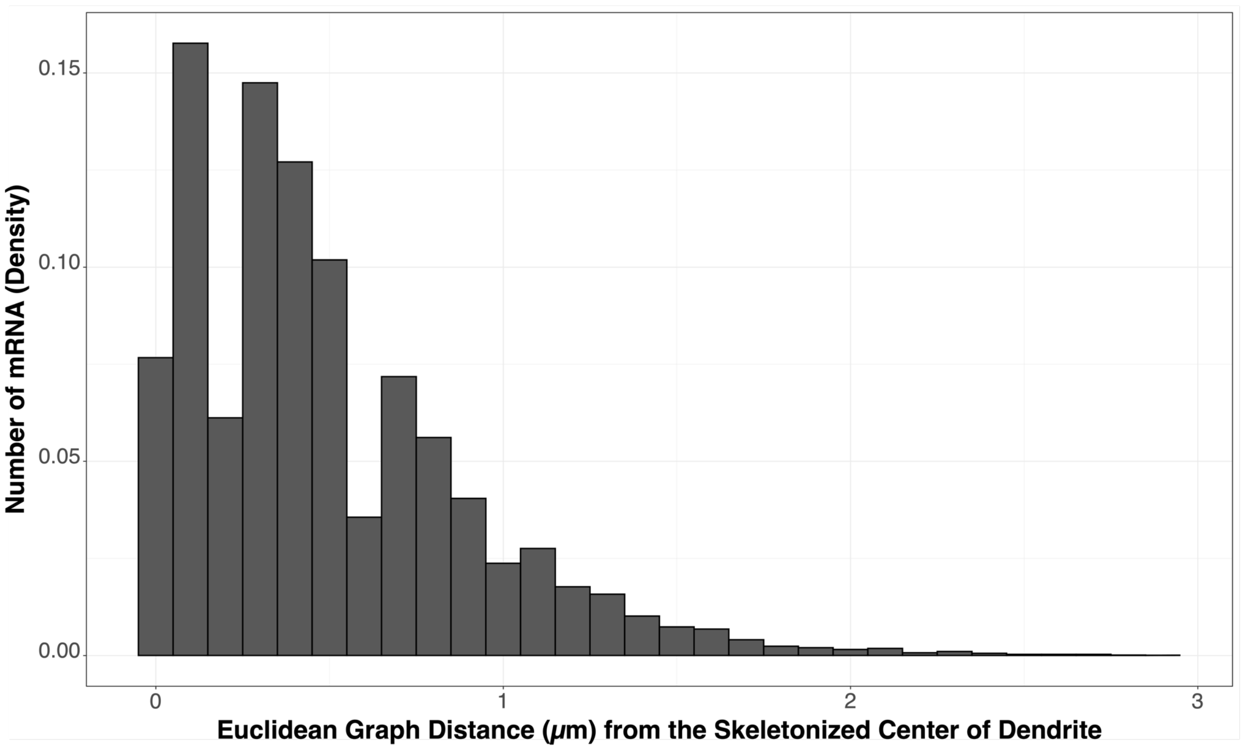


### Supplementary Fig. 4

a

b

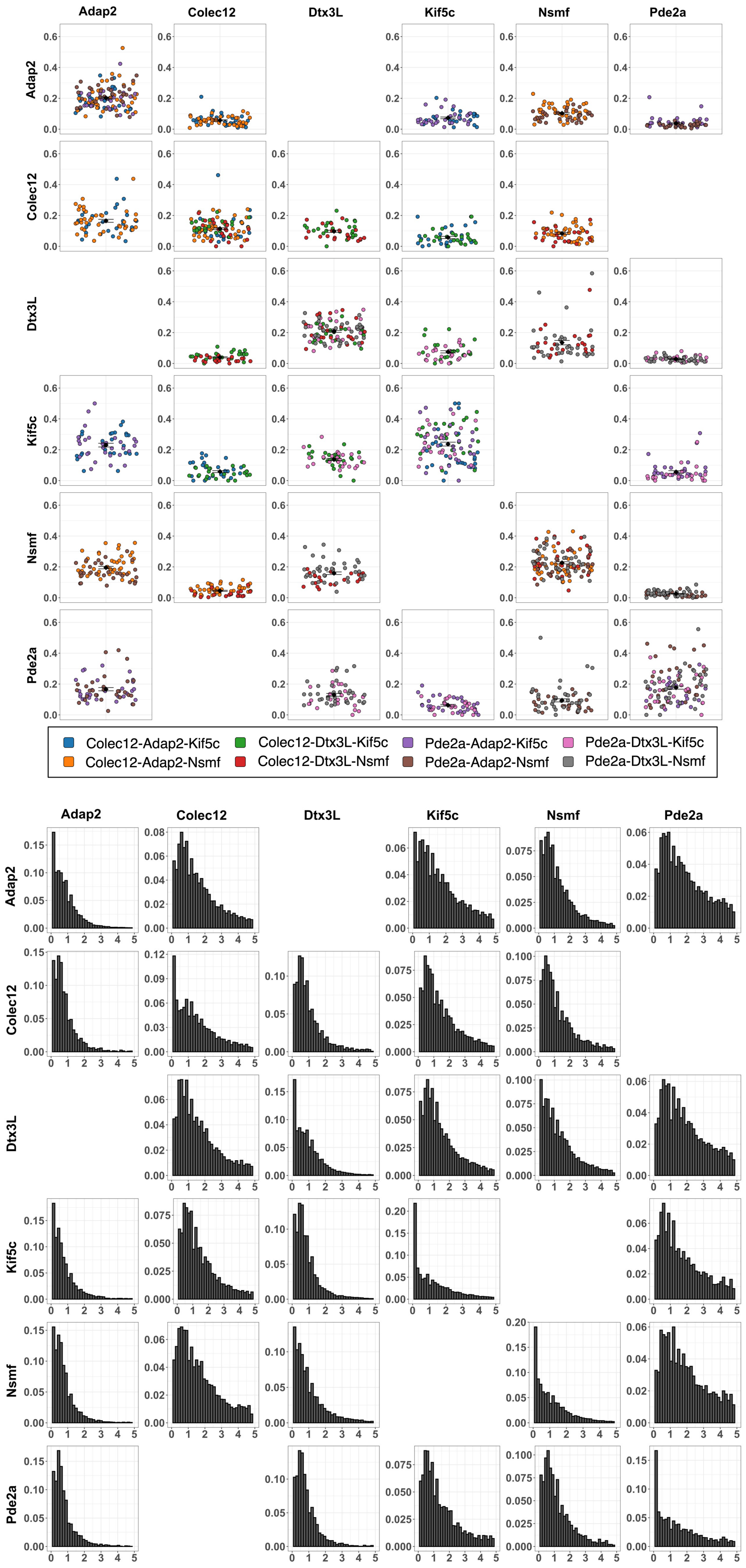


### Supplementary Fig. 5
