## Supplementary Tables for "Characterizing the Spatial Distribution of Dendritic RNA at Single Molecule Resolution"

| Cy3 (Gene) | mCherry (Gene) | Cy5 (Gene) | Number of Neurons Imaged | Number of Neurons Analyzed (after QC) |
|---|---|---|---|---|
| *Colec12* | *Adap2* | *Kif5c* | 43 | 32 |
| *Colec12* | *Adap2* | *Nsmf* | 64 | 49 |
| *Colec12* | *Dtx3L* | *Kif5c* | 58 | 39 |
| *Colec12* | *Dtx3L* | *Nsmf* | 33 | 27 |
| *Pde2a* | *Adap2* | *Kif5c* | 60 | 49 |
| *Pde2a* | *Adap2* | *Nsmf* | 53 | 42 |
| *Pde2a* | *Dtx3L* | *Kif5c* | 40 | 32 |
| *Pde2a* | *Dtx3L* | *Nsmf* | 59 | 50 |

**Supplementary Table 1:** Summary of the gene probes used in each color channel and numbers of neurons imaged/analyzed per triplet probe combination.

| | ***Colec12-Adap2-Kif5c*** | ***Colec12-Adap2-Nsmf*** | ***Colec12-Dtx3L-Kif5c*** | ***Colec12-Dtx3L-Nsmf*** | ***Pde2a-Adap2-Kif5c*** | ***Pde2a-Adap2-Nsmf*** | ***Pde2a-Dtx3L-Kif5c*** | ***Pde2a-Dtx3L-Nsmf*** |
|---|---|---|---|---|---|---|---|---|
| ***Adap2*** | 3.59e-03 ± 9.05e-04 | 5.36e-03 ± 1.43e-03 | | | 4.90e-03 ± 1.23e-03 | 5.23e-03 ± 1.35e-03 | | |
| ***Colec12*** | 1.23e-03 ± 5.47e-04 | 1.62e-03 ± 4.61e-04 | 1.79e-03 ± 8.71e-04 | 7.46e-04 ± 3.27e-04 | | | | |
| ***Dtx3L*** | | | 3.95e-03 ± 1.71e-03 | 2.29e-03 ± 6.35e-04 | | | 4.71e-03 ± 9.06e-04 | 4.51e-03 ± 1.00e-03 |
| ***Kif5c*** | 1.17e-03 ± 6.14e-04 | | 1.92e-03 ± 9.50e-04 | | 1.47e-03 ± 7.74e-04 | | 2.26e-03 ± 1.19e-03 | |
| ***Nsmf*** | | 2.64e-03 ± 9.12e-04 | | 2.42e-03 ± 1.14e-03 | | 2.68e-03 ± 1.03e-03 | | 2.89e-03 ± 1.44e-03 |
| ***Pde2a*** | | | | | 1.17e-03 ± 4.50e-04 | 7.40e-04 ± 3.13e-04 | 1.17e-03 ± 2.71e-04 | 8.06e-04 ± 3.74e-04 |

**Supplementary Table 2:** Mean mRNA densities (number of detected mRNAs per pixel area of neuronal compartment) and standard deviations in the dendrites across the triplet probe combinations. Refer to **Figure 1A**.

| | *Colec12-Adap2-Kif5c* | *Colec12-Adap2-Nsmf* | *Colec12-Dtx3L-Kif5c* | *Colec12-Dtx3L-Nsmf* | *Pde2a-Adap2-Kif5c* | *Pde2a-Adap2-Nsmf* | *Pde2a-Dtx3L-Kif5c* | *Pde2a-Dtx3L-Nsmf* |
|---|---|---|---|---|---|---|---|---|
| ***Adap2*** | 8.18e-03 ± 1.78e-03 | 1.47e-02 ± 2.70e-03 | | | 1.79e-02 ± 4.79e-03 | 1.58e-02 ± 2.67e-03 | | |
| ***Colec12*** | 3.20e-03 ± 1.50e-03 | 4.98e-03 ± 1.50e-03 | 4.34e-03 ± 1.91e-03 | 2.20e-03 ± 7.50e-04 | | | | |
| ***Dtx3L*** | | | 1.17e-02 ± 4.28e-03 | 7.45e-03 ± 2.22e-03 | | | 1.13e-02 ± 2.85e-03 | 7.63e-03 ± 1.83e-03 |
| ***Kif5c*** | 2.03e-03 ± 9.92e-04 | | 4.77e-03 ± 2.53e-03 | | 3.42e-03 ± 1.69e-03 | | 4.16e-03 ± 1.76e-03 | |
| ***Nsmf*** | | 4.80e-03 ± 1.99e-03 | | 5.32e-03 ± 2.71e-03 | | 5.71e-03 ± 1.99e-03 | | 4.96e-03 ± 2.84e-03 |
| ***Pde2a*** | | | | | 5.60e-03 ± 2.05e-03 | 4.96e-03 ± 2.45e-03 | 5.52e-03 ± 2.38e-03 | 3.59e-03 ± 1.97e-03 |

**Supplementary Table 3:** Mean mRNA densities (number of detected mRNAs per pixel area of neuronal compartment) and standard deviations in the soma across the triplet probe combinations. Refer to **Figure 1A**.

| Gene | Mean Graph Distance (μm) from Soma |
|---|---|
| *Adap2* | 34.0877818 |
| *Colec12* | 35.2664164 |
| *Dtx3L* | 35.6162905 |
| *Kif5c* | 35.3362085 |
| *Nsmf* | 33.5015466 |
| *Pde2a* | 30.0922761 |

**Supplementary Table 4:** Localized transcripts' mean graph distance from the soma calculated per gene.
